# Material-Driven Fibronectin Assembly Rescues Matrix Defects due to Mutations in Collagen IV in Fibroblasts

**DOI:** 10.1101/2020.01.06.895839

**Authors:** Elie Ngandu Mpoyi, Marco Cantini, Yuan Yan Sin, Lauren Fleming, Dennis W. Zhou, Mercedes Costell, Yinhui Lu, Karl Kadler, Andrés J. García, Tom Van Agtmael, Manuel Salmeron-Sanchez

## Abstract

Basement membranes (BMs) provide structural support to tissues and influence cell signaling. Mutations in COL4A1/COL4A2, a major BM component, cause eye, kidney and cerebrovascular disease, including stroke. Common variants in these genes are risk factors for intracerebral hemorrhage in the general population. However, the contribution of the matrix to the disease mechanism(s) and its effects on the biology of cells harboring a collagen IV mutation remain poorly understood. To shed light on this, we engineered controlled microenvironments using polymer biointerfaces coated with ECM proteins laminin or fibronectin (FN), to investigate the cellular phenotype of primary fibroblasts harboring a *COL4A2^+/G702D^* mutation. FN nanonetworks assembled on poly(ethyl acrylate) (PEA) induced increased deposition and assembly of collagen IV in *COL4A2^+/G702D^* cells, which was associated with reduced ER size and enhanced levels of protein chaperones such as BIP, suggesting increased protein folding capacity of cells. FN nanonetworks on PEA also partially rescued the reduced stiffness of the deposited matrix and cells, and enhanced cell adhesion through β_1_-mediated signaling and actin-myosin contractility, effectively rescuing some of the cellular phenotypes associated with *COL4A1/4A2* mutations. Collectively, these results suggest that biomaterials are able to shape the matrix and cellular phenotype of the *COL4A2^+/G702D^* mutation in patient-derived cells.

## 1. Introduction

Basement membranes (BMs) are specialized extracellular matrices (ECM) structures that provide structural support to tissues as well as influence cell behavior and signaling.^[1–3]^ Major components are laminins, collagen IV, perlecan and nidogen.^[2, 4]^ The vertebrate genome encodes six collagen IV alpha chains, α1(IV)-α6(IV), encoded by the *genes COL4A1-COL4A6* that generate three collagen IV networks, namely α1α1α2(IV), α3α4α5(IV) and α5α5α6(IV).^[3, 5]^

Mutations in COL4A1 and COL4A2 result in a rare familial multi-systemic disease encompassing vascular, eye, kidney and muscle defects.^[3]^ The COL4A1 cerebrovascular disease affects the small vessels of the brain and can result in intracerebral hemorrhage, porencephaly (the formation of cerebral cavity due to intracerebral hemorrhage), and white matter hyperintensities.^[6–11]^ The vast majority of mutations affect glycine residues of the collagen repeats which alter protein structure.^[5, 12–14]^ Interestingly, common variants within these genes have been identified as risk factors for intracerebral hemorrhage,^[10]^ myocardial infarct,^[15]^ coronary artery disease,^[16]^ and vascular stiffness.^[17]^ Thus, collagen IV is emerging as a major component for vascular diseases.^[7, 9, 10, 15–22]^ For many of these diseases, there is an urgent need for novel treatments, which will require a more detailed understanding of the underlying molecular mechanisms.

*COL4A1/4A2* mutations such as the *COL4A2^+/G702D^* mutation, a classical glycine mutation that substitutes a glycine residue for aspartic acid,^[8]^ results in matrix defects in patients and mouse models, possibly due to reduced levels of collagen IV caused by intracellular retention in the endoplasmic reticulum (ER).^[8, 14, 23–26]^ The ER is the site where three alpha chains interact to form a triple-helical collagen IV protomer which after secretion generates a collagen IV network in the matrix. ER retention of mutant protein can cause ER stress, which can activate the unfolded protein response (UPR), a homeostatic response that can become pathogenic when chronic.^[24]^ Whereas the pathogenic mechanisms of collagen IV mutations remain unclear, a combination of matrix defects and ER stress likely contribute to pathogenesis.

Recent data indicate that at least some of the cellular and mouse phenotypes due to *COL4A1/4A2* mutations can be modulated. Targeting ER stress using the FDA-approved chemical chaperone phenyl butyric acid, (PBA) decreased intracellular collagen IV retention and ER stress, and increased collagen IV secretion and deposition in the BM in mice, associated with reduced cerebrovascular disease.^[8, 27–29]^ However, PBA did not ameliorate the renal defects,^[27]^ with recent evidence from mouse models supporting cell and/or tissue-specific disease mechanisms whereby the contribution of ER stress and matrix defects to disease may differ.^[13]^ This has led to the suggestion that the matrix defects also need to be targeted in order to develop treatments that address the phenotypes in different tissues.^[27]^ However, the effects of *COL4A1/4A2* mutations on BM function and characteristics remain poorly understood and, importantly, the effects of the matrix on the behavior and cellular phenotype of *COL4A1/4A2* mutant cells remains unknown. Addressing these gaps in our knowledge could identify potential therapeutic targets and/or approaches.

Matrix composition, surface topography and the physical properties of the substrate influence the behavior of cells, and engineered biomaterials provide a powerful approach to modulate the microenvironment of cells and investigate their behavior and function in response to this altered environment.^[30]^ In tissues, the ECM is a critical part of the cell environment, and fibronectin (FN) is a major ECM component that binds other matrix proteins, growth factors and cell surface receptors such as integrins, which are also collagen receptors. The biological activity of FN can be controlled by adsorbing it onto polymers with defined chemistry, such as poly(ethyl acrylate) (PEA) and poly(methyl acrylate) (PMA). Adsorption onto PEA results in physiological-like FN nanonetworks that provide better availability of cell and growth factors binding regions, while PMA leads to the formation of globular aggregates, Figure 1.^[31–36]^ The ability to adsorb matrix components onto PEA and PMA, therefore, provides a powerful system to present the cell with controlled distribution and conformation of matrix proteins. For FN, the physical and chemical similarity of PEA and PMA provides a robust system to investigate the effect of the FN network as a model of an altered ECM on cells expressing mutant collagen _IV._^[36]^

**Figure 1.**
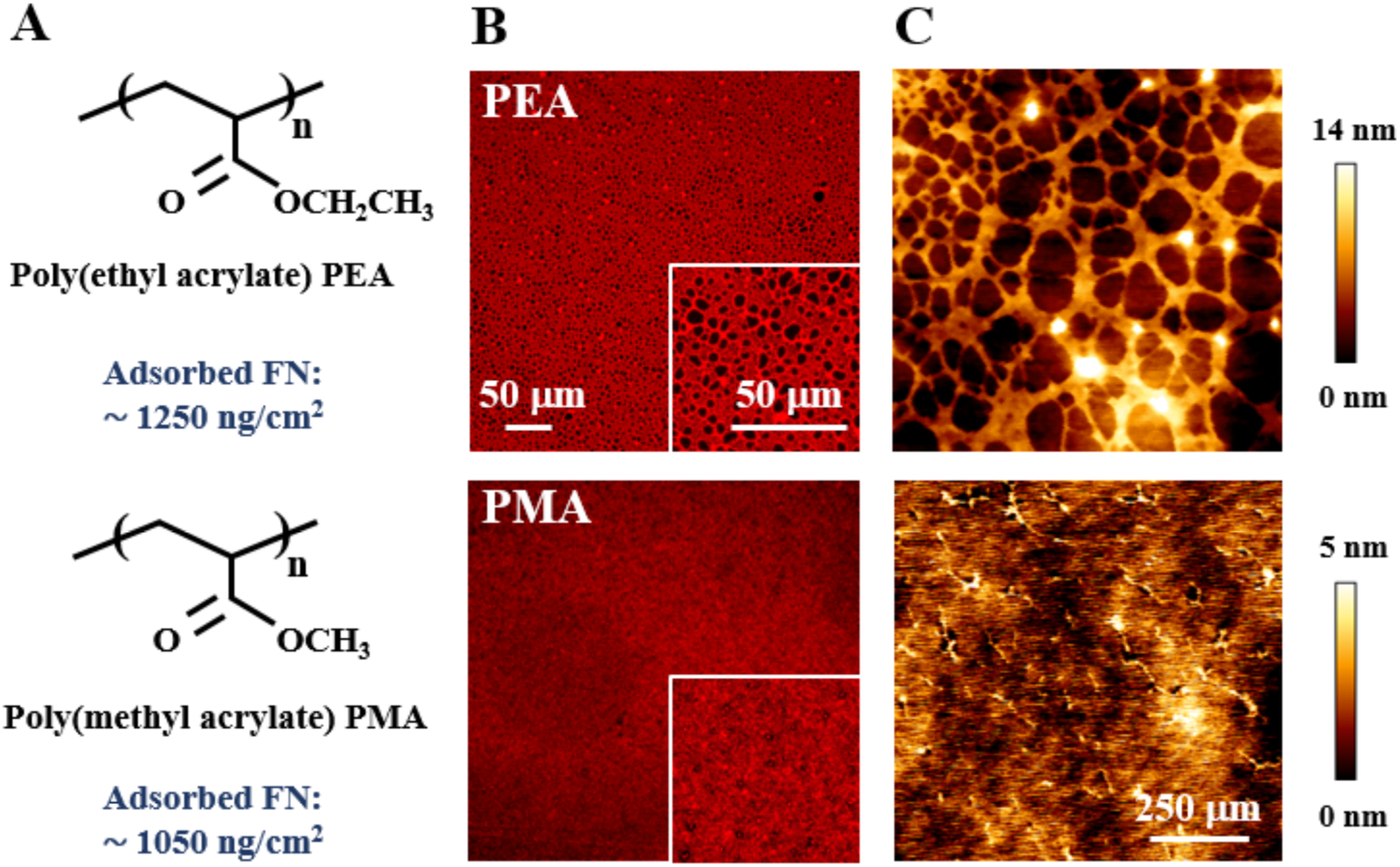
Molecular organization of FN after adsorption on PEA and PMA polymer substrates. (A) Structure of PEA and PMA, and amount of adsorbed FN from a solution concentration of 20 µg/mL for 1 hour. (B) Immunostaining of FN. Insert: high magnification images. Scale bars: 50 µm. (C) AFM height images of FN.

Here, we have uncovered that FN nanonetworks assembled on PEA promote the deposition of collagen IV in primary patient cells carrying a *COL4A2^+/G702D^* mutation. This is associated with an increase in the protein folding capacity of the cell, ameliorated stiffness of the cells and their matrix, and increased cell adhesion through focal adhesions. We show that these effects of FN are via specific integrin-mediated signaling. These data indicate that biomaterials can offer a controlled matrix to modulate the cellular phenotype of collagen IV mutations.

## 2. Results

### 2.1. Col4 deposition and ER-stress of *COL4A2^+/G702D^* fibroblasts

*COL4A2^+/G702D^* fibroblasts (MT) were cultured on fibronectin (FN)-coated substrates, which control the organization and presentation of adsorbed FN, and on glass controls (uncoated). Adsorbed FN assembled into physiological-like nanonetworks on PEA, whilst it retained a globular conformation on PMA, with similar FN amounts adsorbed, Figure 1. To investigate the effects of the controlled matrix on the behavior of COL4A2 mutant cells, we assessed COL4A2 deposition. Image analysis revealed reduced COL4A2 deposition on glass after 1 and 7 days (Figure 2A-C) in MT cells compared to WT, as previously reported.^[8, 45]^ To provide the first insight into the secreted collagen IV network of COL4A1 or COL4A2 mutant cells, we also assessed the fractal dimension of the collagen IV network as a descriptor of its interconnected fibrillar organization (Figure 2E). The collagen IV network of *COL4A2^+/G702D^* fibroblasts on glass is characterized by reduced fractal dimension and a disrupted fibrillar organization; this confirms the “structural” effect of the glycine mutations on the collagen IV network. This was accompanied by a slower growth rate for the MT cells,^[6]^ which were less confluent, with cell size often enlarged and formation of patchy populations indicative of cell death compared to WT cells (Figure S1).

**Figure 2.**
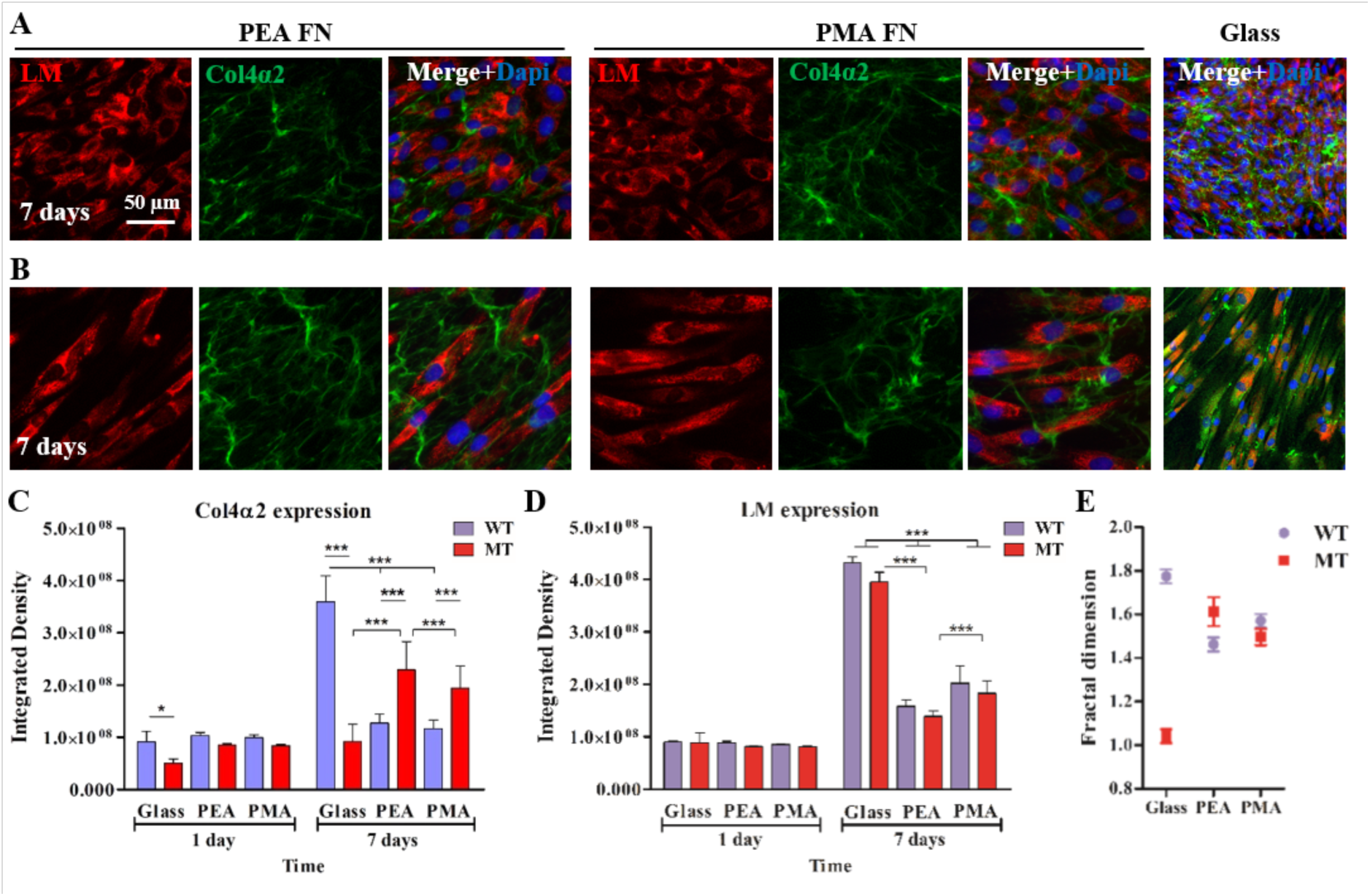
Deposition of Col4A2 (green) and LM (red) by control (A) and mutant fibroblasts (B) on PEA and PMA coated with FN. Cells were grown on PEA and PMA substrates-coated with FN 20 µg/ml and on glass for 2 h under serum free conditions; then with serum before fixation at different time points (1 and 7 days). Cells were also simultaneously stained with DAPI (Blue). Scale bars: 50 µm (for all micrographs). Quantification of expressed Col4A2 (C) and LM (D) using integrated density per cell. Fractal dimension analysis of secreted Col4A2 (E). Data presented as mean ± SD, N ≥10; and analyzed with an ANOVA test; *p < 0.05; ***p<0.001. Important statistical significance differences between the WT and the MT cells are indicated, including between the substrates for the MT. WT, wild type control fibroblast cells; MT, COL4A2^+/G702D^ mutant fibroblast cells.

Strikingly, we observed that MT fibroblasts on FN-coated polymers displayed increased deposition of secreted collagen IV, especially after 7 days of culture (Figure 2C). This was particularly significant for cells on the FN nanonetworks on PEA, where collagen IV deposition by MT cells was the highest and the fractal feature of the collagen meshwork was recovered (Figure 2E). This enhanced deposition on FN-coated PEA compared to FN-coated PMA (where FN remains globular) and glass was confirmed by in-cell-western (Figure S2A-B), ELISA (Figure S2C) and fluorescent staining without cell permeabilization to avoid detecting intracellular proteins (Figure S3). It was also independent on the presence of serum in the culture medium (Figure S4). Increased matrix deposition was also apparent in electron microscopy images, which showed enhanced fibrillar matrix deposition in MT cells on PEA compared to PMA (Figure S5). Moreover, this response appeared specific for FN-coated PEA, as LM-coated samples did not yield the same effect (Figure S6). The increased deposition did not extend to other major BM components as LM protein levels decreased on FN-coated polymers (Figure 2D), and was independent of increased bulk secretion (Figure S7).

We next investigated whether the higher collagen IV levels on FN-coated PEA were accompanied by a reduction in intracellular collagen IV retention. Co-staining was performed with protein disulphide isomerase (PDI), a marker of the endoplasmic reticulum (ER), as ER stress is associated with an increase in ER size and area of the cells.^[8]^ This revealed significantly lower levels for MT cells on PEA-FN compared to MT cells on glass and PMA-FN, suggesting a decrease in ER area in MT cells on FN-coated PEA (Figure 3A and B). Swollen vesicles were also more apparent by electron microscopy in cells cultured on PMA compared to PEA (Figure S5).

**Figure 3.**
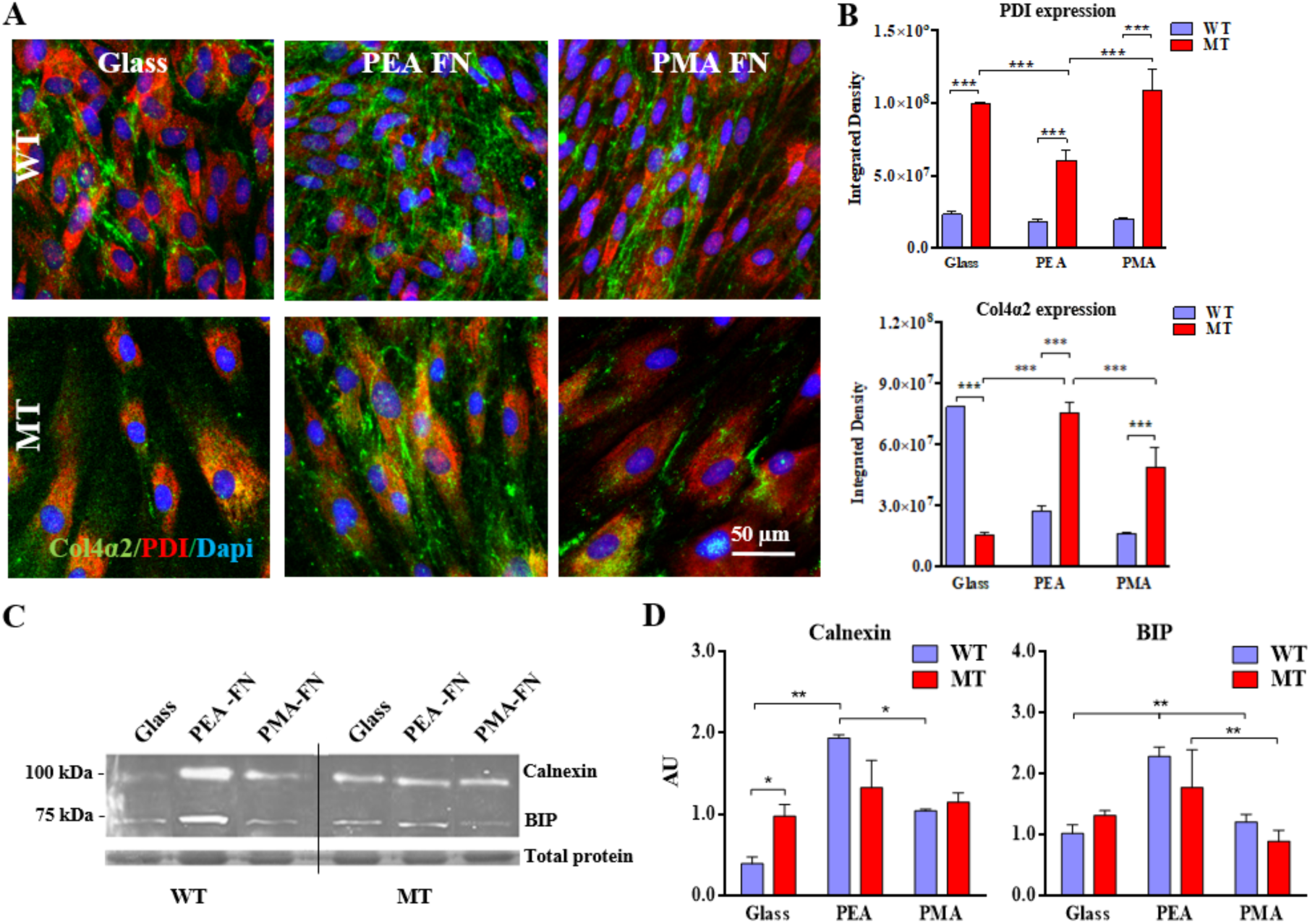
Protein folding capacity of *COL4A2^+/G702D^* fibroblast cells. Staining of COL4A2 (green) and PDI (red) in WT (top row) and MT fibroblasts (bottom row), also simultaneously stained with DAPI (Blue). Cells were cultured on FN-coated PEA and PMA substrates for 7 days. Scale bars: 50 µm (for all micrographs) (A). Quantification of expressed PDI and Col4A2 integrated density measurements per cell (B). Western blot analysis of ER stress markers Calnexin (90 kDa) and BIP (78 kDa) levels in cell lysates from WT and MT cells cultured for 7 day (C); Densitometry of western blots shown in arbitrary units (AU) (D). Data presented as mean ± SD, N =4; *p<0.05, **p<0.01, ***p<0.001; N-number: 12. WT, wild type; MT, *COL4A2^+/G702D^* fibroblasts.

The protein folding capacity of the cells is determined by the levels and activities of the protein folding machinery including chaperones such as BIP; for example, levels of BIP are associated with levels of protein secretion in yeast.^[46]^ To investigate whether the decrease in ER area in the MT cells on FN-coated PEA was associated with increased protein folding capacity, the protein levels of chaperones BIP and calnexin were measured by western blot (Figure 3C). Interestingly, this revealed elevated BIP protein levels in both cell lines when grown on PEA-FN compared to PMA, indicating a mutation-independent effect. This suggests that PEA-FN may lead to an increase in the protein folding capacity of the cells.

### 2.2. Elastic properties of *COL4A2^+/G702D^* fibroblasts and of their ECM

The characteristics of the matrix influences cell behavior and cell function including those of vascular cells.^[47]^ However, the effect of any collagen IV mutation on the characteristics of the basement membrane remains unclear. To address this, the effect of the *COL4A2^+/G702D^* mutation on the biomechanical properties of the cells and their matrix was analyzed via atomic force microscopy (AFM) by measuring the overall stiffness of matrix and cells, and then of the decellularised ECM only. The Young’s modulus of the MT cells, measured via nano-indentation using beaded cantilevers, was 10-fold lower than that of WT cells after 7 days of culture on glass, PEA-FN or PMA-FN, Figure 4A. Values ranged mainly from 500 to 1000 Pa on glass, from 2000 to 5000 Pa on PEA-FN, and from 500 to 1500 Pa on PMA-FN for WT cells (Figure S8A), and from 10 to 50 Pa on glass, from 100 to 350 Pa on PEA-FN, and from 25 to 150 Pa on PMA-FN for MT cells (Figure S8B). In any case, culture on PEA-FN increased the Young’s modulus of cells and their matrix (Figure 4A).

**Figure 4.**
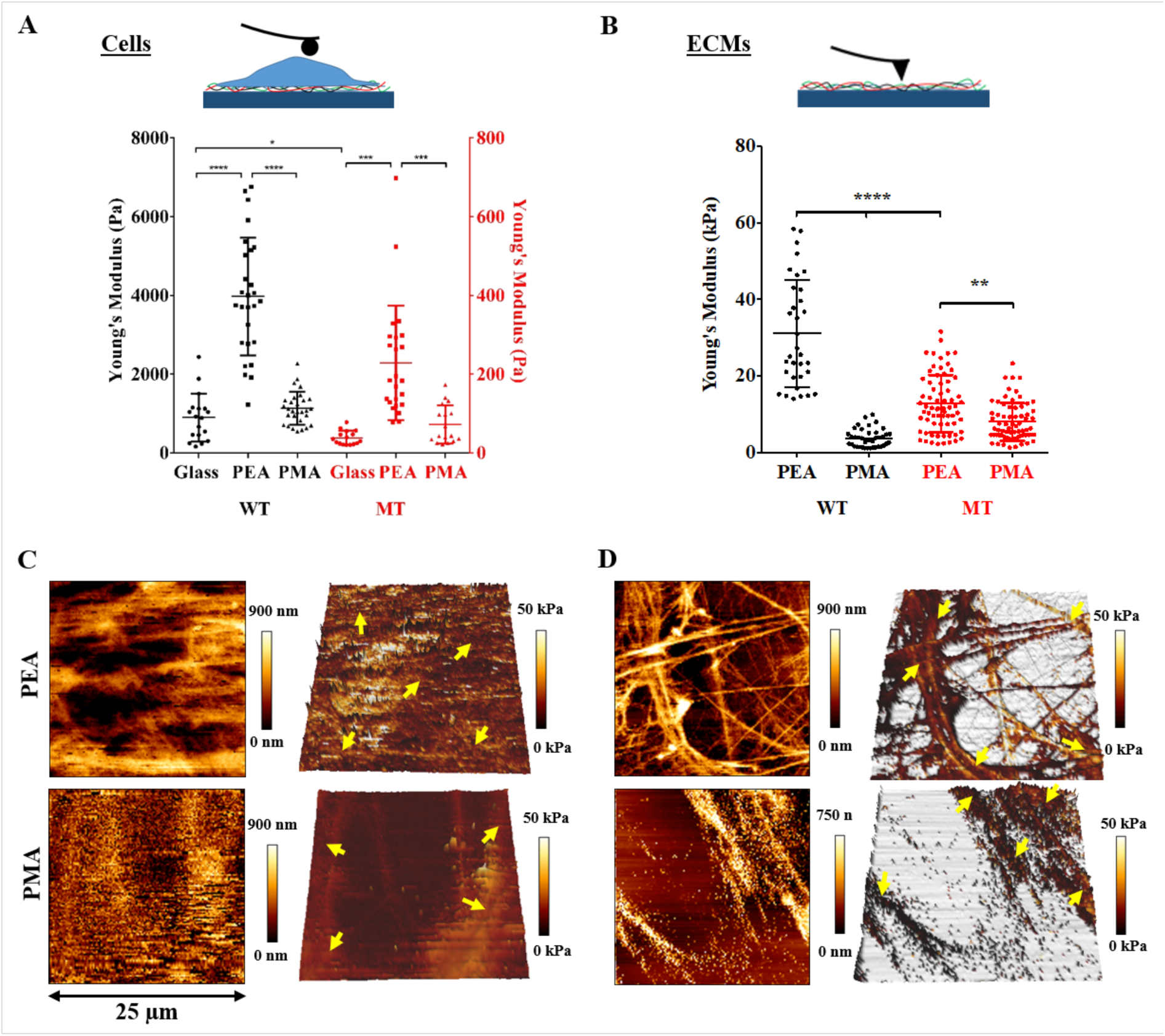
Elastic modulus analysis of mutant cells and secreted ECMs. Young’s modulus of WT and MT cells on the different substrates measured via AFM force mapping; cells were cultured for 7 days and then indented with a cantilever mounted with a 4.83 µm silica bead (A). Young’s modulus of ECMs obtained using the Hertz model on at least 20 measurements (N ≥20) taken from the points indicated by the yellow arrows in the AFM images C and D (B). AFM height images (first column) and 3D reconstruction (second column) of ECMs after decellularization of WT (C) and MT (D) cells; cells were cultured on FN-coated PEA and PMA for 7 days and then decellularized using 20 mM ammonium hydroxide (NH_4_OH) solution, leaving the ECMs intact; then, AFM quantitative imaging was carried out in DPBS using a pyramidal tip. The color scale of the 3D reconstruction represents the local Young’s modulus of the ECMs, calculated using the Hertz model. All data are presented as mean ± SD, and analyzed with an ANOVA test; **p<0.01, ***p<0.001, ****p<0.0001. WT, wild type; MT, *COL4A2^+/G702D^* cells.

Most interestingly, the measurement of the elastic properties of the decellularized matrix (Figure 4B) using a pyramidal tip in quantitative imaging mode revealed a partial recover of matrix stiffness when MT cells were cultured on PEA. Indeed, the values ranged from 7.5 to 18 kPa for the MT-secreted matrix, and from 20 to 43 kPa for WT. On PMA-FN the values were instead generally lower than 10 kPa for both cell types. These data suggest that the increased deposition by culture on PEA-FN is able to partially rescue the mechanical properties of the secreted ECM. Moreover, 3D reconstruction of the decellularised ECMs by MT cells cultured on PEA-FN show a clear fibrillar organization (Figure 4C-D). No measurements were made for ECMs secreted by cells on glass, as they often detached from the substrate during the measurements.

### 2.3. Cell adhesion

To gain insights into underlying mechanisms of the altered stiffness and behavior of the MT cells when cultured on FN-coated PEA, we analyzed the early cell adhesion to the substrates, focusing on the development of focal adhesion (FA) complexes and on the role of the cytoskeleton. Focal adhesions were visualized using antibody staining against paxillin, Figure 5A-B. Image analysis revealed increased cell size for MT cells compared to WT (Figure 5C), and well-defined FA plaques for both cell types, mainly located at the cell periphery (Figure 5A-B). To explore the role of contractility, cells were incubated with blebbistatin, which inhibits myosin-II-specific ATPase activity. Treatment with blebbistatin did not affect cell morphology, except a few cell protrusions of MT cells on PEA-FN and PMA-FN (Figure S9A-B).

**Figure 5.**
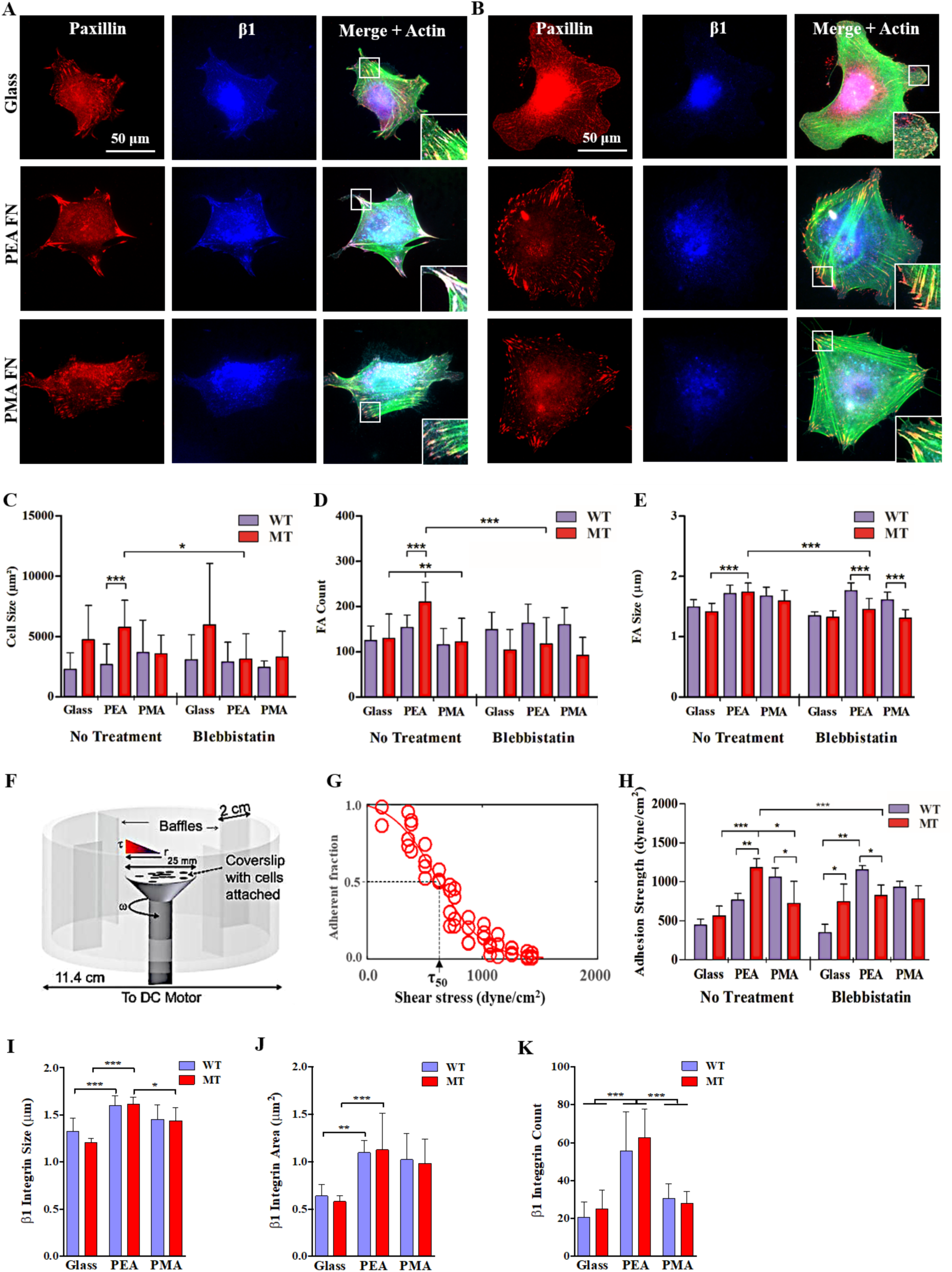
Focal adhesion assembly, β_1_ integrin and cell adhesion strength measurements on FN coated PEA and PMA substrates. Immunofluorescence images of focal adhesions and β_1_ integrin of WT (A) and MT (B) fibroblast cells on glass, and on PEA and PMA coated with 20 µg/ml of FN, for 2h in media under serum-free conditions without or with blebbistatin; cells were fixed and then stained for paxillin (red), β_1_ integrin (blue) and actin (green). Further images are provided in supporting data. Cell size of WT and MT cells (C); number of FAs per cell (D); size of FAs (E). Schematic of spinning disk assay (F). Characteristic detachment profile (G) showing fraction of adherent cells (f) as a function of surface shear stress (τ). Adhesion strength measurements for WT and MT cells (H). Quantification of integrated β_1_ integrin size (I) area (J) and count (K) performed with ImageJ and an N =3. Data presented as mean ± SD, N ≥12, and analyzed with an ANOVA test; *p<0.05, **p<0.01, ***p<0.001. Important statistical significance differences between the WT and the MT cells are indicated, including between the substrates for the MT. WT, wild type; MT, *COL4A2^+/G702D^* fibroblasts.

To quantify the maturation level of the FAs on the different surfaces, FA count and size (defined as the length of the major axis of the FA plaque) were analyzed (process detailed in Figure S10). The number and size of FAs for MT cells were significantly higher on PEA-FN compared to glass and PMA-FN (Figure 5D and E), whilst no differences were found for WT cells. Strikingly, treatment with blebbistatin only affected the MT cells cultured on PEA-FN by reducing the number and size of the FAs to the same levels as the other surfaces (Figure 5D and E), indicating myosin II-regulated adhesion.^[48]^ It is also worth noting that ligand availability, as defined by the concentration of the FN coating solution, affected MT cell behavior, in terms of cell size and FA number only on PEA-FN whilst it had no effect for MT cells cultured on PMA-FN (Figure S9D and E).

The increased size and number of FAs for MT cells on PEA-FN correlated with higher adhesion strength, measured using a spinning disk hydrodynamic shear assay, Figure 5F.^[39, 49, 50]^ Detachment profiles (adherent fraction f versus shear stress τ) were fitted to a sigmoid curve to obtain the shear stress for 50% detachment (τ_50_), which is defined as the adhesion strength (Figure 5G).^[51]^ MT cells showed statistically higher adhesion strength on PEA-FN compared to glass and PMA-FN. This was reduced when contractility was inhibited using blebbistatin (Figure 5H, Figure S11), in accordance with FA analyses results (Figure 5D and E) and supporting the role for myosin II-regulated adhesion on PEA-FN. Studies at increasing FN coating solution concentrations confirmed a direct relationship between ligand density and adhesion strength, which increased for PEA-FN (Figure S9G and S12).^[39, 50, 51]^ The role of cell cytoskeleton in adhesion to PEA was also confirmed via electron microscopy, which showed enhanced microfilament organization within cells cultured on PEA-FN compared to PMA-FN (Figure S7).

Considering that cell adhesion to FN is mainly mediated by α_5_β_1_ and α_V_β_3_ integrins,^[52]^ we explored their role in the stronger adhesion of MT cells to the FN nanonetworks on PEA compared to PMA-FN or glass. Both integrins were expressed by both cell types adhering on FN-coated substrates (Figure 5A-B and Figure S13). In particular, β_1_ staining showed well-pronounced clusters resembling FA contacts for the WT cells on all the surfaces. In contrast, it was rather dispersed throughout the MT cells (Figure 5A-B), while α_v_ showed a diffused staining in the entire cell on all the substrates for both MT and WT cells (Figure S13A-B). Quantification of integrin staining showed that number, size and area of β_1_ integrin clusters were generally higher for MT and WT cells on PEA-FN than on PMA-FN and glass. For α_v_, on the other hand, the expression levels by MT cells were maintained independently of the substrates (Figure S13C). Collectively, these results indicate higher β_1_ expression by MT cells on PEA-FN, suggesting the involvement of the FN fibrillar network on PEA in cell adhesion and signaling.

As myosin II is important for the FA recruitment of focal adhesion kinase (FAK),^[53]^ we explored the potential role of FAK via Western Blotting and immunofluorescent staining. Well-developed Y397-pFAK clusters were observed on FN-coated polymers, while poorly developed clusters occurred on glass (Figure S14A). Western blotting confirmed higher levels of pFAK for the MT cells on FN-coated substrates compared to glass. The MT cells level of pFAK was elevated on PEA-FN compared to the WT cells (Figure S14B-C), correlating with the observations of the immunostaining. This revealed enhanced FAK signaling on MT cells from FN networks on PEA; note that previous studies have shown little difference for pFAK between FN-PEA and FN-PMA.^[34, 35]^

Besides integrin receptors, the disk-shaped receptor tyrosine kinases discoidin domain receptors (DDRs)^[54]^ also bind collagen.^[55]^ DDR1 is a tyrosine kinase transmembrane receptor that binds collagen IV and other collagens and regulates cell adhesion, differentiation, migration and proliferation.^[54, 55]^ As the effects of collagen IV mutations on DDR1 expression are unknown, we performed immunostaining which revealed DDR1 expression on the membrane surface for both cell types on all surfaces (Figure S14D). Intriguingly, western blotting detected higher DDR1 protein levels for the MT cells on glass compared to the FN-coated substrates, and statistically higher than WT cells (Figure S14E-F). This suggests an inverse correlation between DDR1 protein levels and amount of collagen IV deposition, whereby reduced collagen deposition due to the COL4A2 mutation leads to upregulation of DDR1.

To further corroborate the implication of integrin binding and cell contractility in the change of the MT cell phenotype when cultured on FN nanonetworks, COL4A2 deposition was assessed on PEA substrates coated with mutated fibronectin molecules or in the presence of blebbistatin. In particular, mouse ΔRGD FN (FN lacking the RGD motif) and mouse syn FN (FN with a mutation on the DRVPPSRN synergy binding site) were coated onto PEA, forming FN nanonetworks lacking the ability to bind both α_5_β_1_ and α_v_β_3_ (ΔRGD FN) or to reinforce binding to α_5_β_1_ (syn FN), respectively (Figure S15A).^[56, 57]^ Immunofluorescent staining confirmed higher deposition and fibrillar organization of collagen IV by MT fibroblasts only occurred on PEA coated with wild type FN, as both mutations reverted the rescue effect of the wild type FN nanonetworks, Figure 6. This suggests a major role for α_5_β_1_ integrin. Interestingly, the unavailability of the cell binding and of the synergy domain on the FN nanonetwork only affected the behavior of MT cells, whilst WT fibroblasts were unaffected (Figure S15B). Moreover, the addition of blebbistatin to the media during the culture also ablated MT cells collagen IV deposition by inhibiting cell contractility, Figure 6.

**Figure 6.**
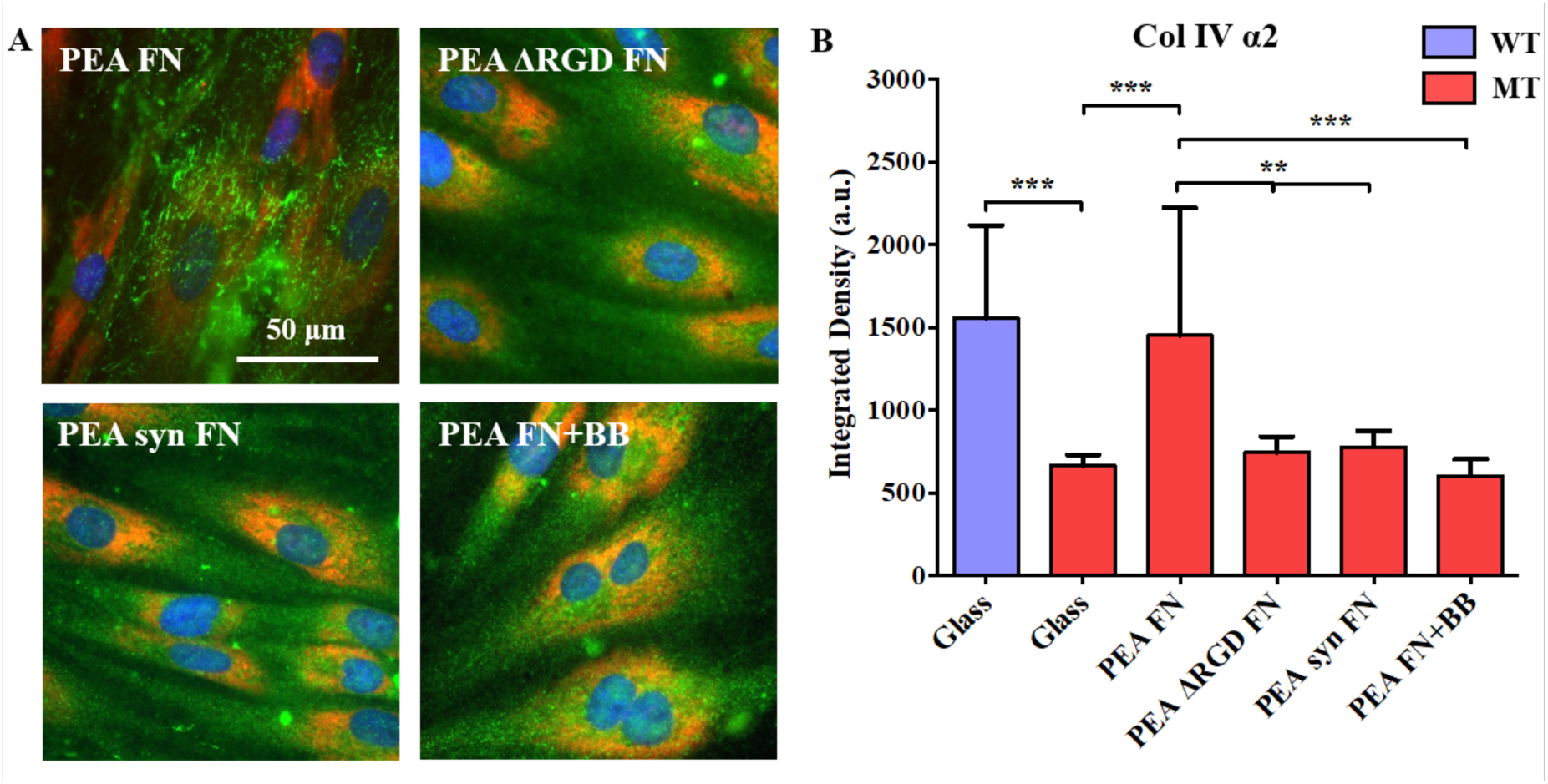
Deposition of COL4A2 by MT fibroblasts on FN nanonetworks is regulated by integrin binding and cell contractility. COL4A2 staining (green) of cells grown on PEA substrates coated with either FN, FN without the RGD domain (ΔRGD FN), FN with a mutation on the synergy binding site (syn FN), or in the presence of blebbistatin (BB) in the culture medium for 7 days; LM (red) and nuclei (blue) were also stained (A). Quantification of the integrated density of COL4A2 staining after variance filtering (B). Controls are WT fibroblasts and MT fibroblasts cultured on glass. Data presented as mean ± SD, N ≥12, and analyzed with an ANOVA test; *p<0.05, **p<0.01, ***p<0.001. WT, wild type; MT, *COL4A2^+/G702D^* fibroblasts.

## 3. Discussion

It is known that material properties, such as stiffness, topography and chemistry, can alter cell phenotype and drive cellular processes such as cell migration, cell signaling and (stem) cell differentiation.^[58–60]^ These processes involve the secretion of new ECM at the cell-material interface, whose composition and nature are modulated by the properties of the biomaterials on which cells grow. Here, we demonstrate that engineered biomaterials have the potential to modulate matrix defects due to mutations in collagen IV in fibroblasts. The *COL4A2^+/G702D^* mutation and other *COL4A1/4A2* mutations result in lower deposition and incorporation of collagen IV into the BM.^[8, 45]^ Here, results show that FN nanonetworks assembled on the surface of PEA increase the deposition of COL4A2 in *COL4A2^+/G702D^* mutant cells, with similar levels to WT (normal) cells. This phenomenon is shown by several approaches, immunofluorescence (including without permeabilization, to rule out the possibility of staining intracellular COL4A2 and after a decellularization assay), in-cell western and ELISA. We also demonstrate reduced ER stress (reduced ER area – PDI) in MT cells triggered by FN-PEA; this is accompanied by a likely increase in protein folding capacity, with increased levels of molecular chaperone BIP, Figure 7.

**Figure 7.**
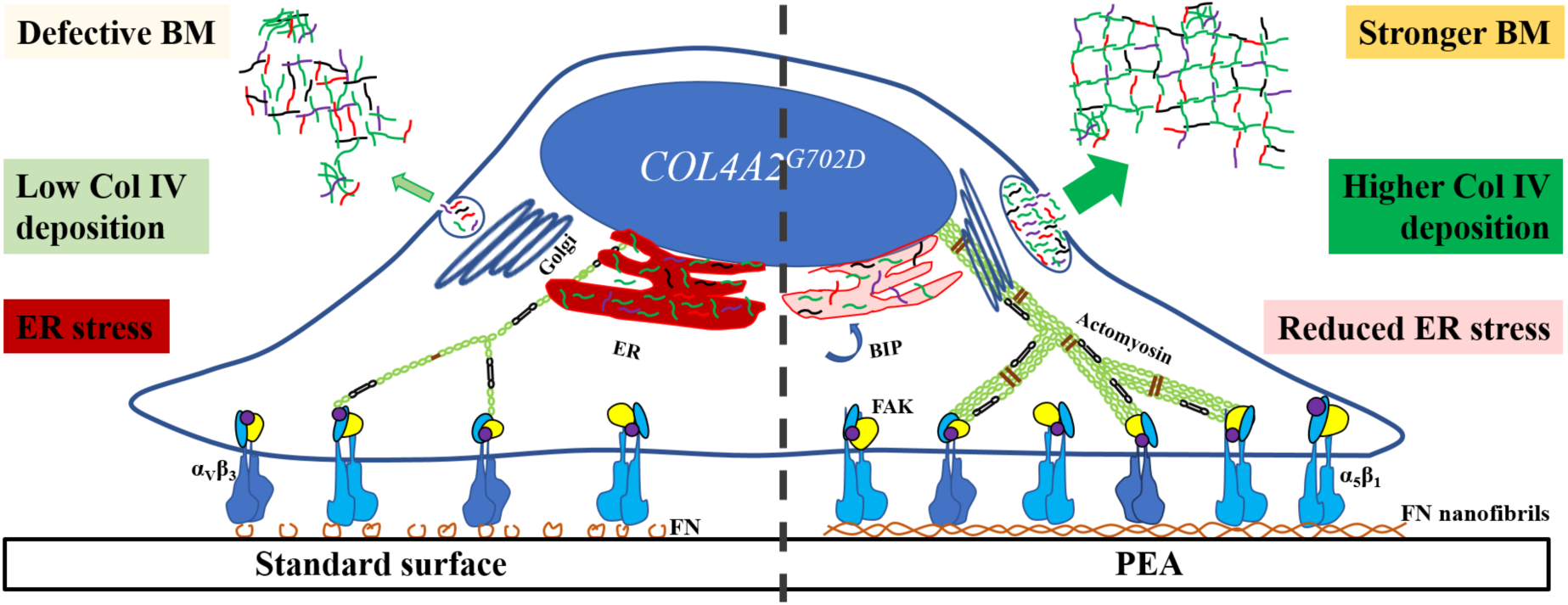
Conceptual scheme. Illustration of the effect of the fibrillar FN nanonetworks on PEA, which induces increased deposition of Col4A2 by *COL4A2^+/G702D^* mutant cells.

MT cells do not increase COL4A2 deposition when cultured on LM interfaces (PEA, PMA, glass) or interfaces where FN remains globular (glass, PMA). The effect is specific to FN nanonetworks assembled on PEA, which appear to enable a partial rescue of the retention and intracellular accumulation of COL4A2 in MT cells. Larger MT cells were found on PEA-FN, with higher FA count and size compared to the cells cultured on PMA-FN or glass. Moreover, these FAs contain higher density of β_1_ integrins, which is the main receptor involved during the initial cell interaction with the material-driven FN network and physiological FN matrices.^[34]^ FN adsorption on PEA unfolds the protein leading to the availability of domains that promote FN-FN interactions, such as FNI_1–5,_ enabling self-assembly into fibrils, recapitulating aspects of cell-mediated FN fibrillogenesis.^[35, 36]^ This material-driven FN matrix also favors enhanced exposure of the integrin-binding region FNIII_9-10_ along with the growth factor binding region FNIII_12-14_.^[31, 33, 60]^ FN assembled on PEA is recognized by β_1_ integrins in *COL4A2^+/G702D^* fibroblasts, Figure 5. Higher expression of β_1_ has been previously correlated to α_5_β_1_ integrin-mediated adhesion to FN matrices, as it happens here for MT cells on FN-PEA, Figure 5.^[51]^ The role of β_1_ in rescuing the deposition of COL4A2 in MT cells is demonstrated by the use of mutant fibronectins, Figure 6. Binding of α_5_β_1_ involves simultaneous availability of the synergy DRVPPSRN and RGD sequences within FN.^[34]^ FNs lacking the RGD peptide or the synergy sequence are also assembled into nanonetworks on PEA (Figure S15A), but they do not lead to enhanced matrix deposition of COL4A2 in MT cells, Figure 6.^[33, 36, 61]^ Interestingly, our data are in agreement with recent *in vivo* data in *C. elegans* supporting the role of integrin in promoting the incorporation of collagen IV into basement membranes from secreted proteins, independently of actual increasing in collagen secretion.^[62]^ Also, we show that the contractile machinery of the cells (ROCK) is activated by binding of β_1_ integrins to FN networks on PEA as the use of blebbistatin reduces adhesion strength only on PEA (Figure 5G), and further ablates the rescue of COL4A2 deposition, Figure 6. This ability of FN-assembled on PEA to trigger cell contractility has been previously demonstrated in cell differentiation processes.^[35]^ Mechanical properties, especially stiffness, of cells and their surrounding ECM play important roles in many biological processes including cell growth, motility, division, differentiation, tissue homeostasis, stem cell differentiation, tumor formation and wound healing.^[63]^ Changes in stiffness of live cells and ECM are often signs of changes in cell physiology or diseases in tissues.^[64]^ Stiffness analysis via AFM showed that the MT cells and their ECMs were 10-fold softer than the WT cells, directly indicating for the first time effects on matrix stiffness due to collagen IV mutations, Figure 4A. However, MT cells were significantly stiffer on FN assembled on PEA, than on glass and FN-coated PMA. Cell stiffness is related to the networks of F-actin and intermediate filaments inside the cells,^[39, 49, 65, 66]^ which are generally observed in a lower amount in the MT than in the WT cells, but not on PEA-FN (Figure 5 and Figure S5). In addition to this, the ECM secreted by MT cells is also stiffer on PEA-FN (Figure 4B), which can be related to the higher amount of deposited COL4A2 on PEA-FN forming interconnected fibrillar networks, Figure 2.^[67]^ It is interesting to note that variants in *COL4A1,* the obligatory protein partner of COL4A2, are genetically associated with vascular stiffness.^[17]^ This study has used patient-derived fibroblasts but can be extended to other type of collagen producing cells in relevant tissues and organs. Analysis into different cell types would include extensive genome editing, as altering collagen IV levels, for example by transfecting expression construct driving the COL4A2 G702D mutation, may cause defects of itself, hampering analysis.

## 4. Conclusion

We show that biomaterials alter the behavior of *COL4A2^+/G702D^* mutant cells by overcoming some of the cellular and matrix defects caused by the mutation. Indeed, physiological-like FN nanonetworks assembled on a specific polymer chemistry (PEA) trigger contractility-dependent strengthening of MT cell adhesion through enhanced recruitment of β_1_ integrins, leading to increased protein folding capacity, increased collagen IV deposition and partial rescue of the mechanical properties of the secreted matrix. Collectively, our results suggest that enhanced integrin signaling, controlled here via biomaterial engineering, influences aspects of the matrix and cellular phenotype of the *COL4A2^+/G702D^* mutation in primary patient cells. These data enhance our understanding of the biological consequences (function/behavior) of *COL4A2* mutations and highlight avenues for potential therapeutic approaches, which are critical to developing personalized therapeutic strategies for intracerebral hemorrhage and other pathologies due to collagen IV mutations.

## 5. Experimental Section

### Preparation of polymer surfaces and protein adsorption

PEA and PMA polymers were synthesized by radical polymerization of ethyl acrylate and methyl acrylate (Sigma, St. Louis, MO), initiated by benzoin at 1 wt% and 0.35 wt%, respectively. PEA and PMA were then dried by vacuum extraction to constant weight and solubilized in toluene 2.5% w/v and 6% w/v respectively. PEA and PMA films (∼1µm) were obtained by spin-casting polymer solutions on glass coverslips (12 or 25mm) at 2000 (PEA) and 3000 (PMA) rpm for 30 seconds, with an acceleration of 3000 rpm/sec. Samples were dried in vacuum for 2 h at 60°C to remove excess toluene. Polymer surfaces were coated with either natural mouse laminin 111 (LM; Invitrogen) or human plasma FN (Sigma) solutions, in Dulbecco’s phosphate-buffered saline (DPBS) at concentrations 2, 10, 20, 50, and 100 µg/mL for 1h and then rinsed with DPBS before use. Mouse plasma fibronectins carrying mutations in the FNIII_10_ module were also used. ΔRGD FN (FN without the RGD sequence) and syn FN (FN with a mutation in the synergy sequence DRVPPSRN>DAVPPSAN) were generated and purified as previously reported.^[37, 38]^ The amount of adsorbed protein was calculated via depletion assay using the bicinchoninic acid working reagent (Thermo Fisher Scientific, Waltham, MA) as previously reported.^[36]^

### Cell culture

Primary dermal fibroblast harboring *COL4A2^+G702D^* mutation (MT) and wild type (WT) fibroblast cells (Tissue Culture Solutions Cell Works, UK) were cultured in DMEM supplemented with 1% Glutamax (Invitrogen™), 1% penicillin and streptomycin solution (Gibco®) (final concentrations of 100 µg/mL), and 10% fetal bovine serum (FBS, Invitrogen™) and kept in an incubator at 37°C with 5% CO_2_. L-ascorbic acid 2-phosphate sesquimagnesium salt hydrate (0.25 mM; Sigma) was administered before cells were fixed to standardize collagen expression and post-translational modifications.^[8]^

### Cell adhesion strength measurements

Cell adhesion strength to adsorbed FN on the polymer surfaces was measured using a spinning disk device.^[39]^ Substrates (25 mm coverslips) were incubated with 2 or 20 µg/mL human FN for 30 min at RT and then incubated with 1% BSA for 30 min at RT. Cells (10,000 cells/cm^2^) were seeded onto substrates and allowed to uniformly attach for several hours (as indicated in figure captions) in media with/without serum in the incubator. Some substrates were treated with 10 µM blebbistatin (B0560-Sigma-Aldrich) (inhibitor of myosin II). Samples were mounted on the spinning disk device, the chamber apparatus was filled with DPBS with 2 mM glucose, and the disk was spun for 5 min at a constant speed with controlled acceleration rates at RT. After spinning, cells were immediately fixed in 3.7% PFA, permeabilized with 1% Triton X-100, and stained with ethidium homodimer (Molecular Probes, Eugene, OR, USA), a DNA-specific fluorescent probe. Cells were counted automatically at specific radial positions using a Nikon TE300 equipped with a Ludl motorized stage, Spot-RT camera, and Image-Pro analysis system, at ×10 magnification. Sixty-one fields (80-100 cells/field before spinning) were analyzed and cell counts were normalized to the number of cells present at the center of the disk, where negligible force is applied. The applied fluid shear stress is given by the formula: 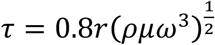, where r is the radial position from the centre of the coverslip, and ρ, µ, and ω are the fluid density, viscosity, and rotational speed, respectively. The fraction of adherent cells (f) was then fitted to a sigmoid curve, Equation 1.

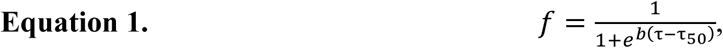

Where b is the inflection slope, and τ_78_ is defined as the shear stress for 50% cell detachment and characterizes the mean adhesion strength of the cell population. The experiment was performed with over eight coverslips per condition and repeated at least twice.

### Immunofluorescence staining

Cells were seeded (5,000 cells/cm^2^) onto PEA and PMA (sterilized under UV for 20 min) coated for 1 h with either wild type or mutant FN (2, 20 µg/mL) or LM (20 µg/mL) and incubated for 2 h in serum-free medium at 37°C, 5% CO_2_, after which the medium was replaced with medium with 10% FBS and 0.25 mM ascorbic acid. Incubation with blebbistatin was performed (if appropriate) and cells fixed with PFA 3.7% for 20 min after specified times (1 and 7 days for matrix secretion), washed 3x with DPBS and permeabilized (0.5% Triton X-100, 10.3 % saccharose, 0.292% NaCl, 0.06% MgCl_2_, and 0.48% HEPES adjusted to pH 7.2) for 5 min and blocked in blocking buffer (BB, 1% Bovine serum albumin, BSA, in DPBS) for 30 min. Cells were incubated with primary antibodies in BB for 1h at RT: monoclonal mouse anti-Col4a2 (Millipore, Cat. No. MAB1910), polyclonal rabbit anti-LM (Sigma, Cat. No. L9393), polyclonal rabbit anti-PDI (Stressgen), polyclonal rabbit anti-FN antibodies, rat anti-mouse CD29 integrin β_1_ (BD Biosciences), human integrin alpha V/CD51 antibody (R&D Systems), mouse vinculin hVIN-1 (Sigma-Aldrich, St. Louis, MO) or polyclonal rabbit paxillin (Thermo Fisher Scientific, Waltham, MA). Cells were washed 3x DPBS/Tween 20 (0.5% w/v), then incubated for 1 h with secondary antibodies. These included Cy™3 AffiniPure goat anti-rabbit (Cat. No. 111-165-003), Cy™3 AffiniPure goat anti-mouse IgG (Cat. No. 115-165-062) (Jackson ImmunoResearch Laboratories Inc), donkey anti-goat Alexa Fluor 568, anti-rat Alexa Fluor 488 (Jackson ImmunoResearch Laboratories), anti-mouse Alexa Fluor 488 (Abcam), anti-rabbit Alexa Fluor 647 (Thermo Fisher Scientific) (in BB). Actin staining was performed using BODIPY FL phalloidin (Invitrogen B607), Alexa Fluor® 350 Phalloidin (Thermo Fisher Scientific) or Rhodamine-Phalloidin (Life Technologies). Samples were washed 3x, then mounted with Vectashield with DAPI to stain the nuclei and visualized using an epifluorescence microscope. Images were taken and channels merged using ImageJ (1.47v).

Fractal dimension analysis of stained secreted collagen IV was carried out using the ImageJ Fractal box count analysis tool, using box sizes of 2, 3, 4, 6, 8, 12, 16, 32, and 64 pixels after thresholding and binarization (Figure S16). Quantification of integrated density was done using ImageJ; the integrated density of each picture was normalized using the number of cells (counted using DAPI) to obtain the integrated density per cell.

Focal adhesions were quantified using the focal adhesion analysis server.^[40]^ Focal complexes, dot-like complexes shorter than 1 µm, were discarded from the analysis.^[41]^ Only isolated cells were used to avoid altered area and roundness values that overlapping cells would have produced. Images were analyzed with ImageJ coupled with an in-house macro processor, and the values of each condition were compared.

### Western blotting

Western blotting was performed as previously reported.^[26]^ Protein extracts were prepared using RIPA buffer containing EDTA protease (Roche Applied Science) and phosphatase inhibitors (Phostop Roche). Membranes were blocked with 5% milk before incubation with primary antibodies calnexin (1/1000, Cell Signaling Technology), BIP (1/40000, BD Transduction), DDR1 (C-20, sc-532, dilution 1:100), polyclonal mouse anti FAK (1:2500 Upstate), and polyclonal rabbit anti p-FAK (Tyr 397) (Merck Millipore). Secondary antibodies used were anti-mouse (1:15000) or anti-rabbit (1:15000) fluorescently tagged antibodies diluted in 50% TBS-T (0.1%) and 50% Seablock with 0.01% SDS; for HRP-conjugated antibodies: donkey monoclonal anti-mouse (1:10000) and donkey monoclonal anti-rabbit antibody (1:10000, GE Healthcare) diluted in 2% BSA-TBST. Protein levels were corrected for Coomassie staining of total protein gels ran in parallel with the western blot gels, and measured using ImageJ software analysis. For analysis of collagen IV secretion, culture medium was switched to serum starved DMEM containing Ascorbic Acid 24 hours before medium and cell collection. Conditioned media was collected and cell lysate were collected following trypsinization after which protein homogenates were prepared in RIPA buffer. Primary antibody used was rat anti-COL4A2 H22 (1:75) from Dr. Tomono Yasuko (Shigei Medical Research Institute) and rabbit anti-GAPDH (1:2000) (Sigma). Statistical analysis was performed on a minimum of three independent experiments using the unpaired t-test.

### Atomic Force microscopy

Atomic force microscopy (AFM) imaging was performed using a JPK Nanowizard® 3 BioScience AFM (JPK Instruments, Berlin, Germany). For imaging of FN adsorbed onto PEA or PMA, samples were rinsed in water, dried gently with a nitrogen flow and imaged in tapping mode using a cantilever with a resonance frequency ∼75 kHz, a spring constant of ∼3 N/m, and a tip radius below 10 nm. The AFM images were 256 × 256 pixels unless specified. Height, phase and amplitude magnitudes were recorded simultaneously for each image. For imaging and force spectroscopy measurements of cells and their secreted matrix, cells were cultured on FN-interfaces for 7 days. All measurements were then taken in liquid (DPBS or CO_2_ independent media) at 37°C. Stiffness of cells and matrix was measured in force mapping mode using a tipless cantilever (Arrow TL1, k = ∼0.03 N/m, NanoWorld, Neuchâtel, Switzerland) mounted with a 4.83 µm diameter silica bead (previously incubated in 1% BSA for 30 min to minimize adhesion). Measurements were performed by indenting living cells on top their nucleus in CO_2_ independent media. Force spectroscopy curves were obtained, after calibration of cantilever sensitivity and spring constant, with a set-point of 5 nN, a z-length of 5 µm, and a constant duration of 1 s. Analyses were performed using the JPK data processing software (v4.3.21) by fitting the force curves with a Hertz model at 500 nm indentation to obtain the Young’s modulus.^[42, 43]^ The stiffness of the ECMs secreted by the cells on the surfaces was also measured via AFM, using the quantitative imaging (QI) mode. To do so, the samples were decellularised to obtain intact matrix deposited by cells via treatment with 20 mM ammonium hydroxide (NH_4_OH) 0.5% (w/v) in warm water until all the cells debris was removed except the insoluble ECMs. The ECMs were then scanned using a pyramidal cantilever (PNP-DB-20, k = ∼ 0.4 N/m, NanoWorld, Neuchâtel, Switzerland). The analysis was performed using the JPK data processing software (v4.3.21) by fitting the force curves with a Hertz model at 50 nm indentation to obtain the Young’s modulus map alongside the contact point height image of the secreted protein matrices. The ECMs were also stained with DAPI to confirm whether the cells were successfully removed.

### Electron microscopy analysis

Cells were cultured on FN-coated PMA and PEA for 7 days and fixed in 2% glutaraldehyde prepared in 100 mM phosphate buffer (pH 7.0). Further processing was performed as previously described.^[44]^

### Statistical analysis

All images were analyzed using ImageJ software (v1.48). The data were statistically analyzed using GraphPad Prism 6 (GraphPad software, La Jolla, CA). Where relevant, one-way or two-way ANOVA tests were performed using a Bonferroni or Tukey post-hoc test to compare all columns, and the differences between groups were considered significant for *p ≤ 0.05, **p ≤ 0.01, ***p ≤ 0.001, and ****p ≤ 0.0001. Error bars represent a standard deviation.

## Supporting Information

The following is the supporting data related to this article: supporting methods and 16 supporting figures. All the original data related to this manuscript are within the depository of the University of Glasgow with https://doi.org/10.5525/gla.researchdata.720.

## Acknowledgements

We would like to thank Dr. Tomono Yasuko (Shigei Medical Research Institute) for the H22 antibody. We thank Rachel Love for the technical support. This work was supported by EPSRC (EP/P001114/1 and EP/F500424/1). MC acknowledges support from MRC through and TVA from Kidney Research UK (RP19/2012). AJG were supported by a US NIH grant (R01 EB024322).

## Author contributions

MSS, TVA and MCantini conceived the research, designed experiments and analyzed data. ENM, MCantini, YYS, LF and YL performed the experiments. AJG and KK designed experiments and edited the manuscript. MCostell provided mutant FNs and edited the manuscript. ENM, MCantini, TVA and MSS wrote the manuscript.

