## Supplementary Materials for "Material-Driven Fibronectin Assembly Rescues Matrix Defects due to Mutations in Collagen IV in Fibroblasts"

### Supporting Methods

*ELISA assay for secreted collagen.* The levels/concentrations of secreted extracellular collagen 4α2(IV) was measured in an indirect ELISA using an ELISA kit in accordance with the manufacturer's instruction (DuoSet R&D Systems). 5,000 cells/cm2 were seeded on 20 µg/mL FN substrates and were maintained in media with FBS for 7 days. Samples were washed and treated with 20 mM ammonium hydroxide to decellularize and get intact ECMs on the surfaces. Then all the samples were blocked with 1% DPBS/BSA for 30 min and incubated with primary monoclonal mouse anti-collagen 4α2(IV) (Millipore, Cat. No. MAB1910) diluted 1:200 in BB for 1h. Triplicate samples were washed with washing buffer then incubated for 1h at RT with HRP conjugated goat anti-mouse antibody diluted in BB. Colour was developed by the addition of the substrate solution (1:1 mix of $H_2O_2$ and tetramethylbenzydine) following a 20-min incubation at RT. At the end of the time, 1/2 volume of stop solution ($H_2SO_4$ 2N) was added. Plates were then read in a Microspectrophotometer, Tecan NanoQuant Infinite M200 Pro plate reader (Männedorf, Switzerland) and the fluorescence were measured at 450 nm and 540 nm. All reagents were provided in the kit.

*In-cell Western assay.* Cells were cultured as above for 7 days, then fixed with 3.7% PFA and then permeabilized with 0.1% Triton X-100 for 5 min, blocked with DPBS/BSA 1% for 30 min for 2h followed by 3 × 10-min washing with 0.1% PBS/Tween 20. Cells were then incubated with primary monoclonal mouse anti-Collagen 4α2(IV) diluted 1:200 in blocking buffer at RT for 1.5h. After 3 × 5-min washing with 0.1% PBST buffer, cells were incubated with infrared-labelled secondary antibody IRDye 800CW (LI-COR, 926-32211) and CellTag 700 Stain (LI-COR, 926-41090) at RT for 1h, followed by 5 × 5-min washing with 0.1% PBST. Substrates were then dried on white paper for infrared signal reading using an Odyssey infrared imaging system.

**Supporting Figures**

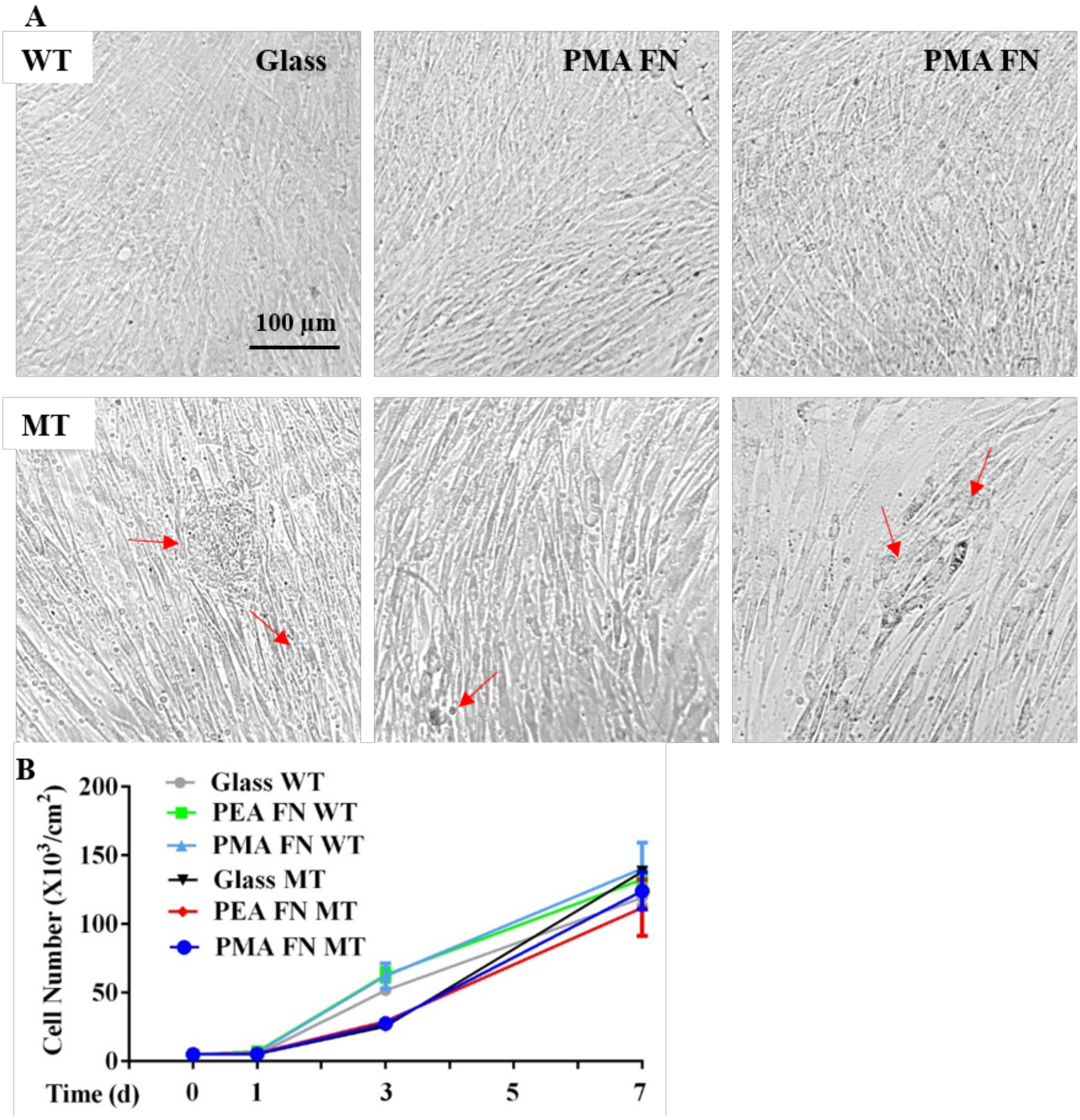


**Figure S1.** Cells morphology and proliferation. A) Light microscopy analysis of the WT and MT fibroblasts of the different substrates at day 7. Red arrows indicate apoptotic MT cells. B) Cells proliferation analysis at the different times: 1, 3 and 7 days. No statistically significant differences found between samples.

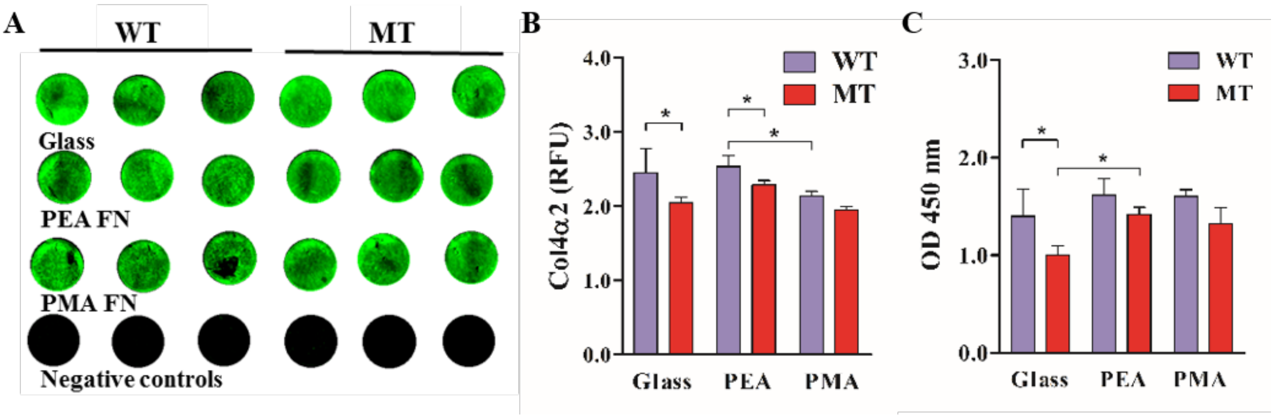


**Figure S2.** Secretion of LM and Col4α2 by control and mutant fibroblasts on PEA and PMA coated with FN. In-cell Western assay for Col4α2 of cells on FN coated substrates after 7 days culture; fluorescence images of the samples (A) and quantifications (B). In addition, an ELISA assay was performed for the deposited Col4α2 after decellularization of cells cultured on FN coated substrates for 7 days culture (C). Negative controls show secondary antibody only and the substrates only. RFU, relative fluorescence units. Only important statistical significance differences between the WT and the MT are indicated, including between the substrates for the MT. WT, wild type; MT, *COL4A2*$^{+/G702D}$ fibroblasts.

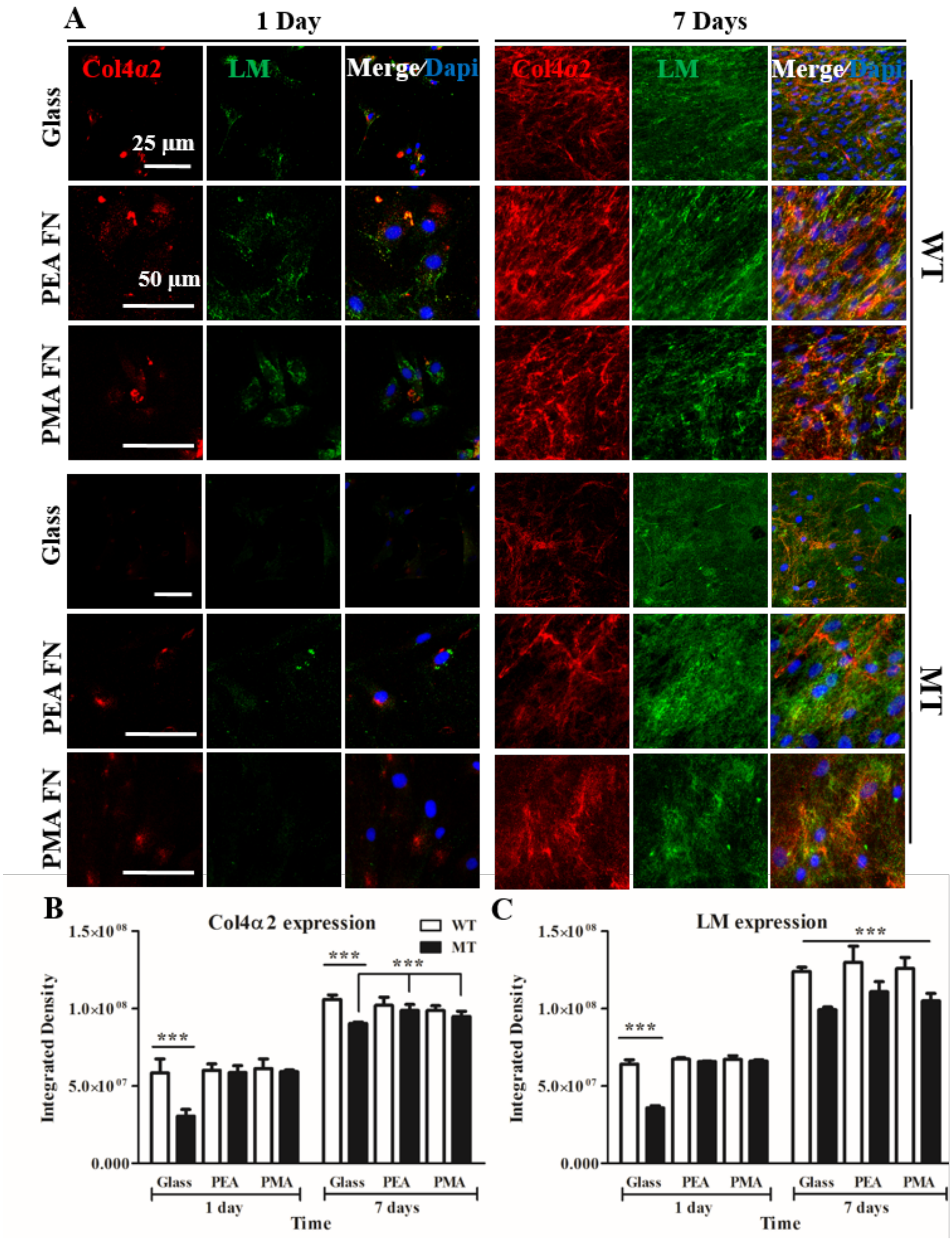


**Figure S3.** Quantification of secreted Col4α2 and LM by control and mutant fibroblasts on PEA and PMA coated with FN with no permeabilization. A) Cells were grown on PEA and PMA substrates-coated with FN 20 µg/ml for 2 h under serum free conditions; then with serum before fixation at different time points (1 and 7 days). Cells were not permeabilized before staining in order to only stain for extracellular proteins. Quantification of expressed Col4α2 (B) and LM (C). Integrated density measurements (of whole image) at different time intervals performed with ImageJ. WT, wild type; MT, COL4A2$^{+/G702D}$ fibroblasts. Data presented as mean ±SD, N ≥10; and analyzed with an ANOVA test; ***p<0.001.

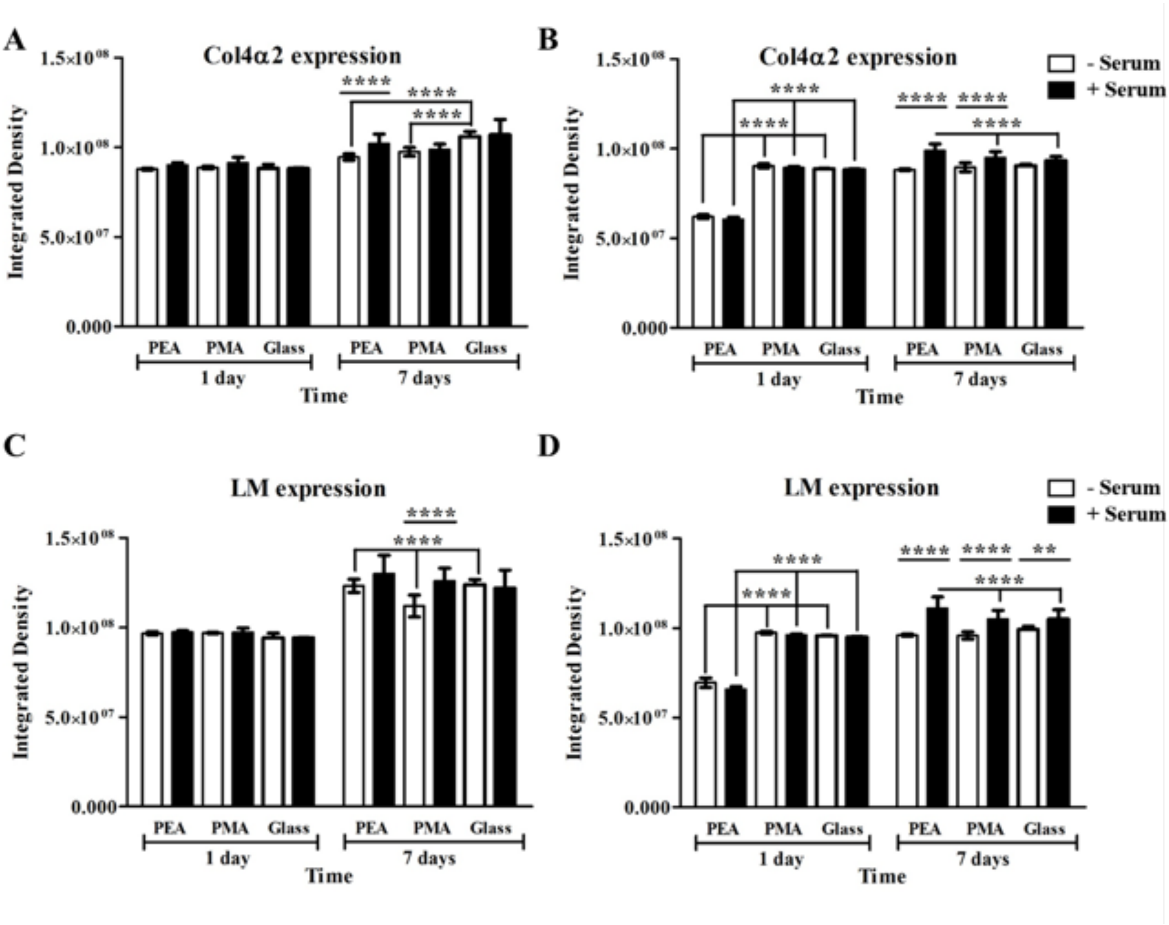


**Figure S4.** Quantification of secreted Col4α2 in mutants cultured with and without serum. Col4α2 and LM in WT fibroblast (A and C) and mutants (B and D) cultured on FN coated polymers for 1 and 7 days with and without serum. Integrated density measurements (of whole image) at different time intervals performed with ImageJ. *p<0.05, **p<0.01, ***p<0.001, ****p<0.0001; N-number: <10.

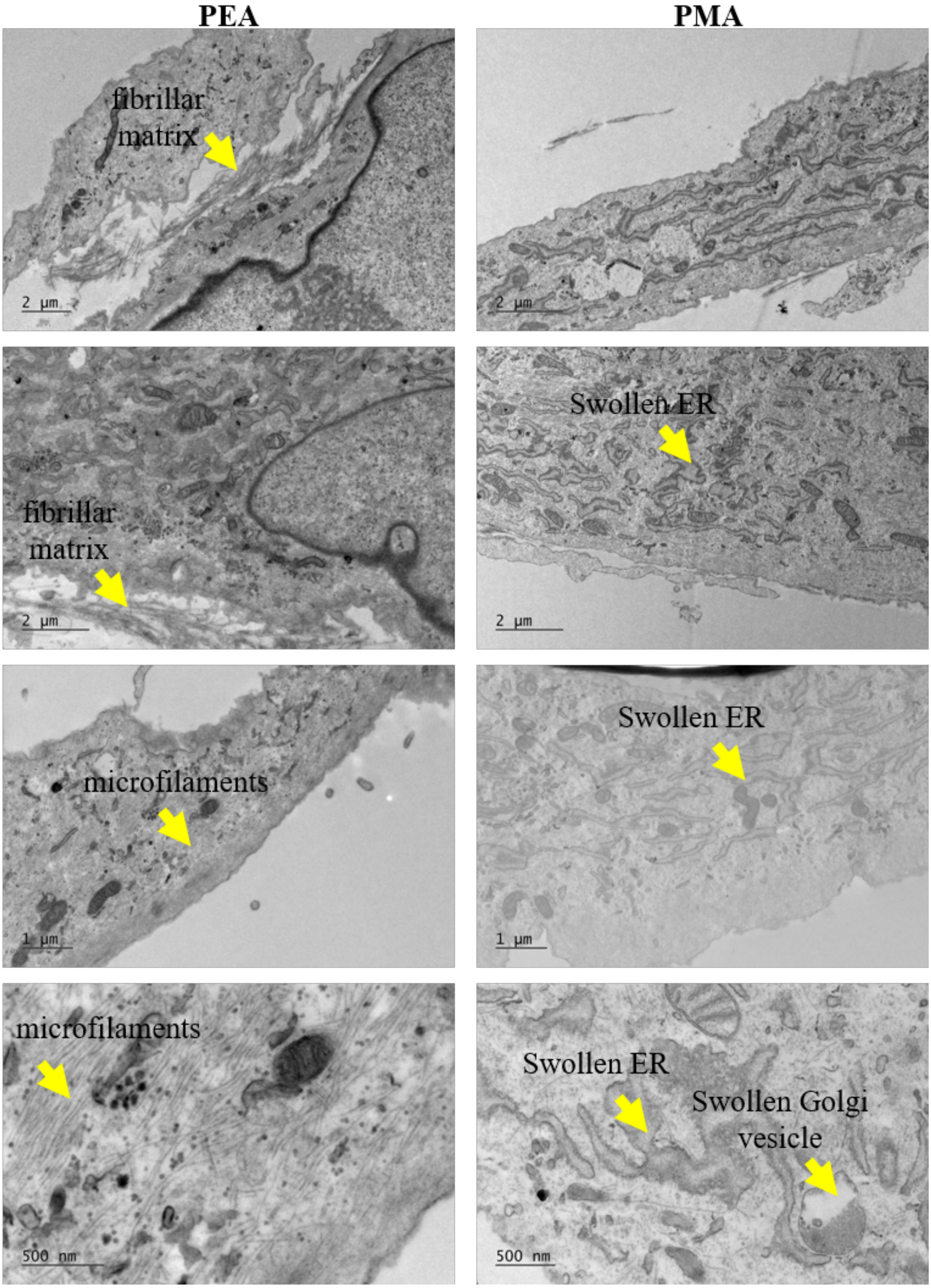


**Figure S5.** Evidence of apparent increased matrix deposition was also apparent in electron microscopy images, which showed enhanced fibrillar matrix deposition in MT cells on PEA compared to PMA. On PMA there were also more apparent vesicles including ER and Golgi vesicles. Cells on PEA displayed more apparent microfilaments. Scale bars shown.

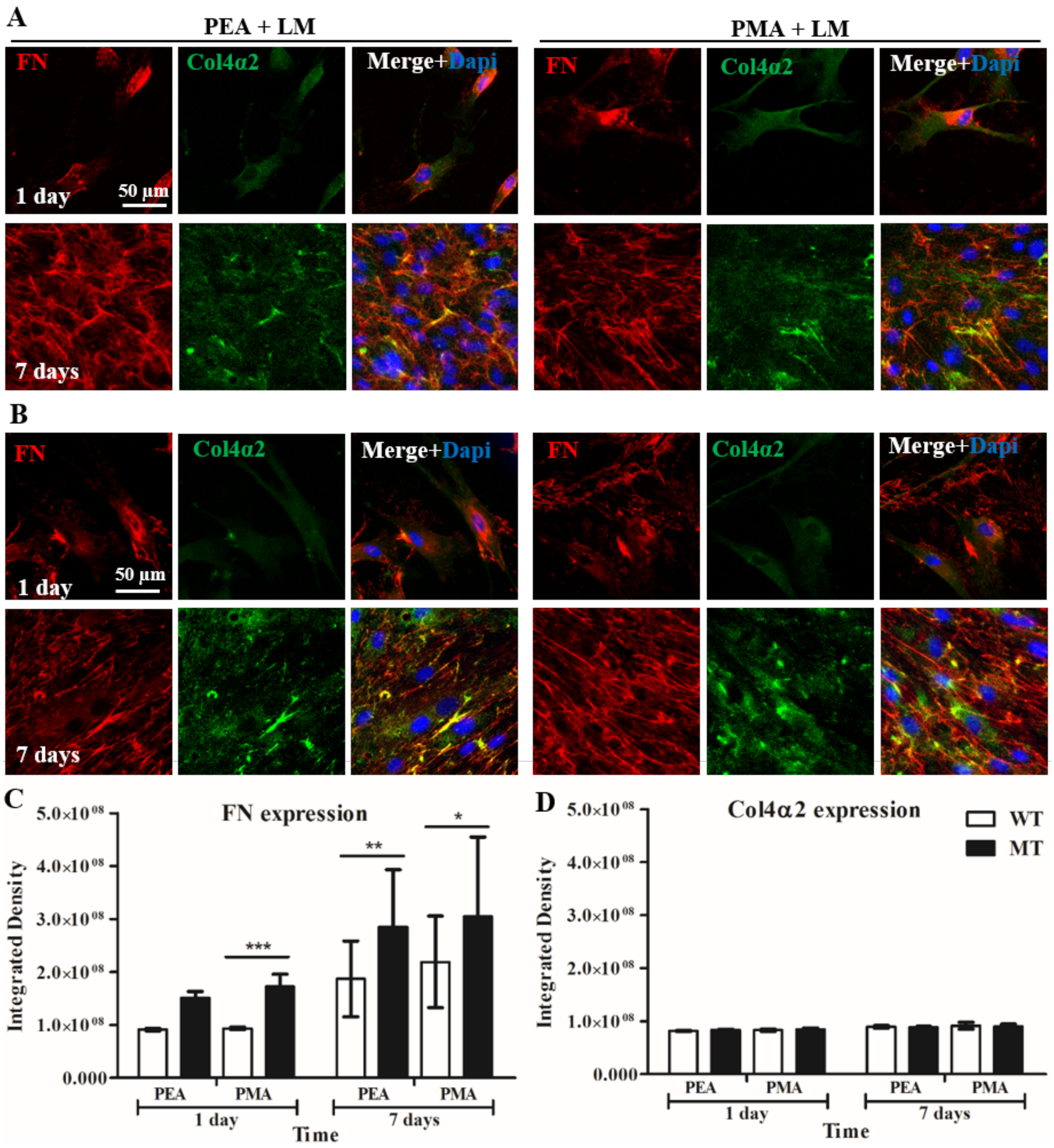


**Figure S6.** Secretion of FN and Col4α2 by control (A) and mutant fibroblasts (B) on PEA and PMA coated with LM. Cells were grown on PEA and PMA substrates-coated with LM 20 µg/ml for 1 and 7 days. Quantification of expressed FN (C) and Col4α2 (D) on LM coated polymers. WT, wild type cells; MT, COL4A2$^{+/G702D}$ fibroblasts. Analyzed with an ANOVA test; *p<0.05, **p<0.01, ***p<0.001. N ≥10.

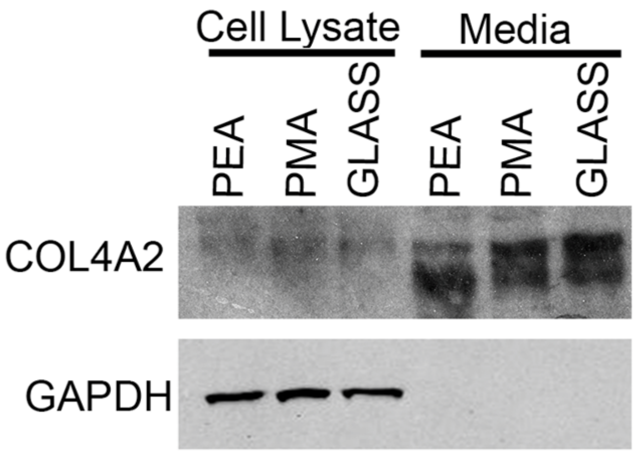


**Figure S7.** Western blot analysis of COL4A2 levels in lysate and media from MT cells cultured for 7 days.

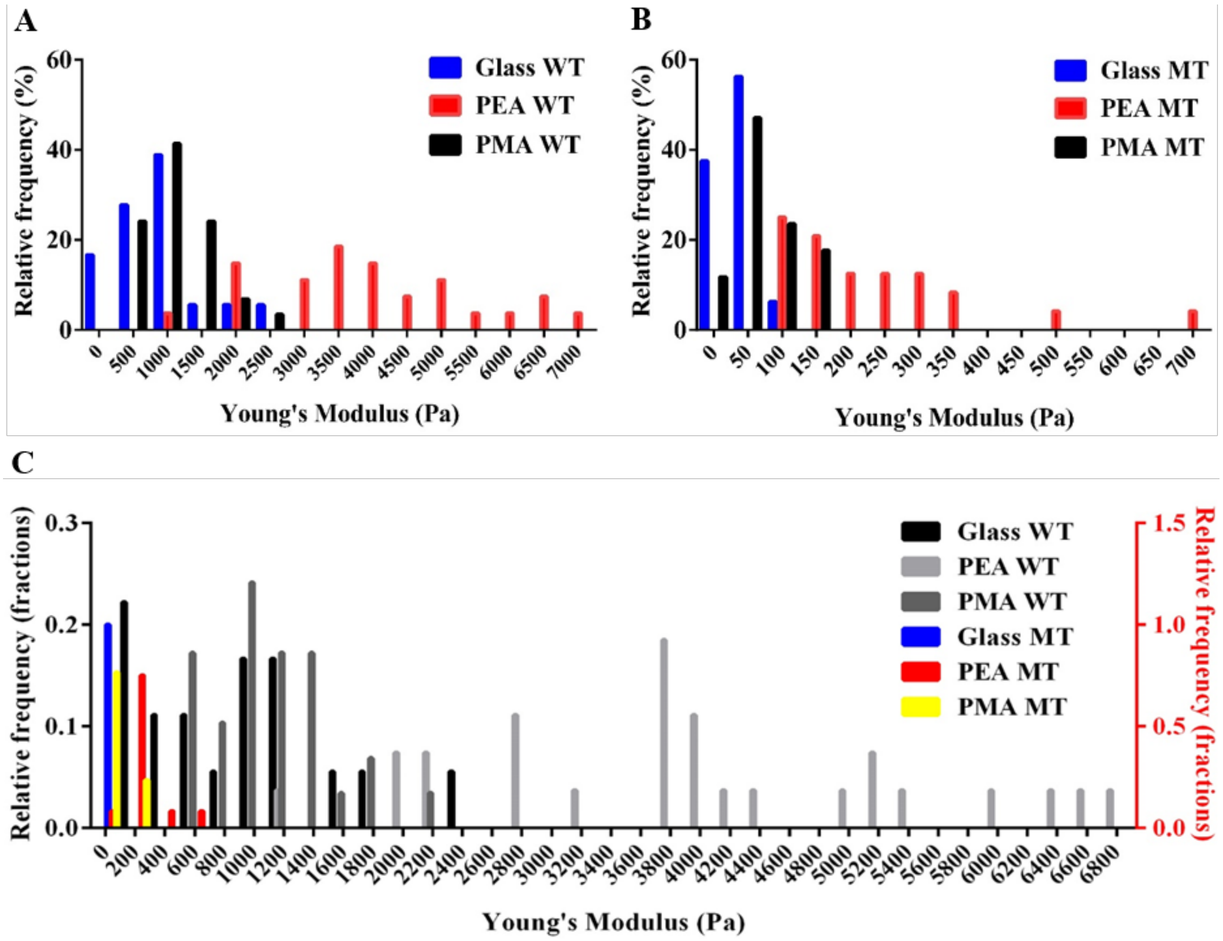


**Figure S8.** Stiffness distribution of WT (A) and MT (B) and combined cell stiffness distributions of WT and MT (C). Cells were cultured on FN coated substrates for 7 days and then scanned live in media using AFM in contact force mapping mode with a tipless cantilever mounted with a 4.85 μm bead. Data presented as mean ±SD, N ≥12; and analyzed with an ANOVA test; *p<0.05, **p<0.01, ***p<0.001, ****p<0.0001; only relevant statistical differences are shown. WT, wild type; MT, mutant cells.

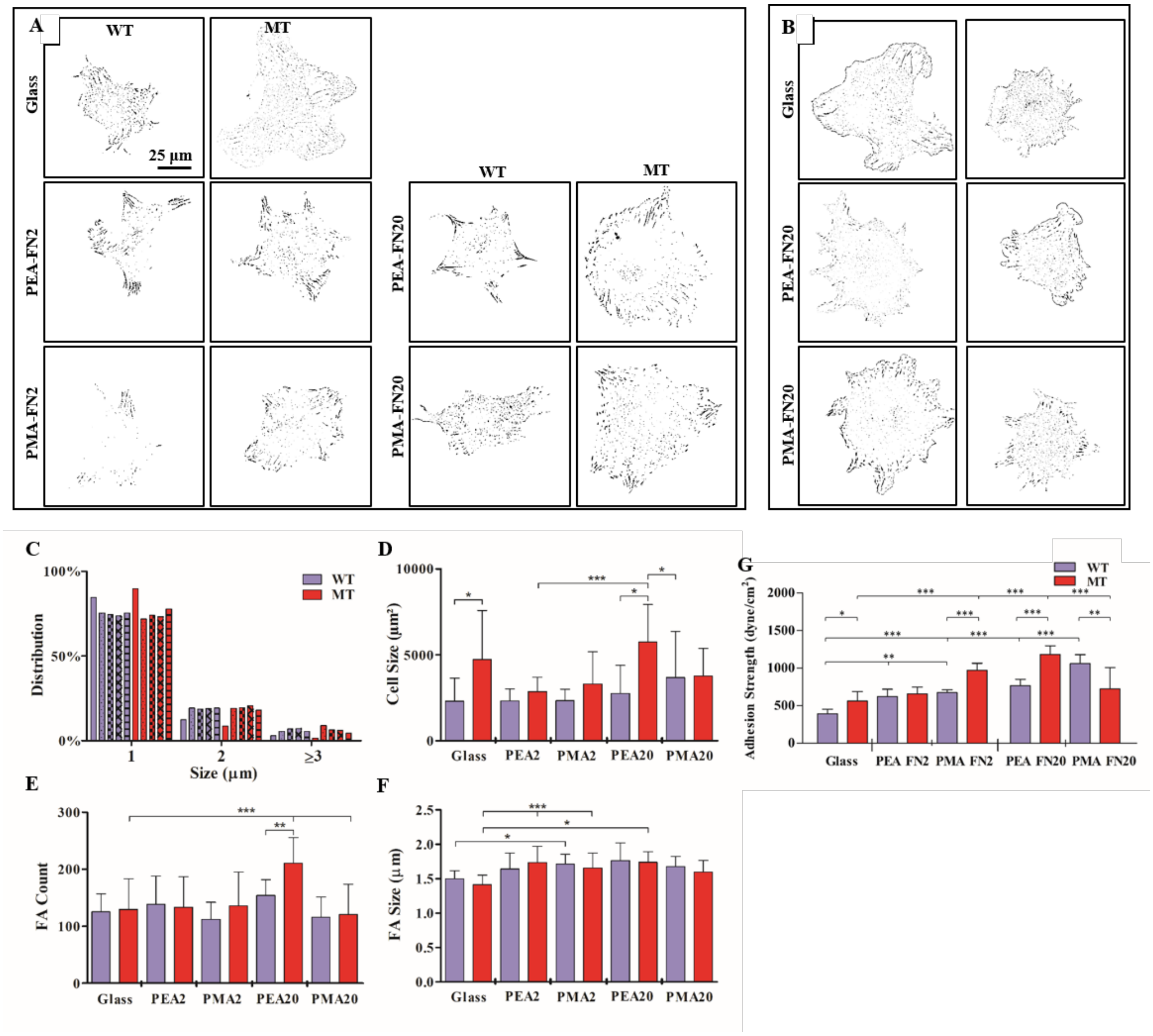


**Figure S9.** Focal adhesion assembly on FN coated PEA and PMA substrates. Representative inverted binary representation of focal adhesions of WT and MT cells on glass and PEA and PMA coated with 2 and 20 µg/ml FN for 2h in media under serum free conditions, fixed and then stained for paxillin (A). Other samples were in media under serum free conditions and treated with blebbistatin (B). Size distribution of focal adhesions in WT and MT Cells (No treatment) (C). Also analyzed were cell size of WT and MT cells (D); number of FAs per cell (E); size of FAs (F). Cell adhesion strength measurements: Influence of fibronectin concentration (G). Data presented as mean ±SD, N ≥12; and analyzed with an ANOVA test; *p<0.05, **p<0.01, ***p<0.001. WT, wild type; MT, *COL4A2*$^{+/G702D}$ fibroblasts.

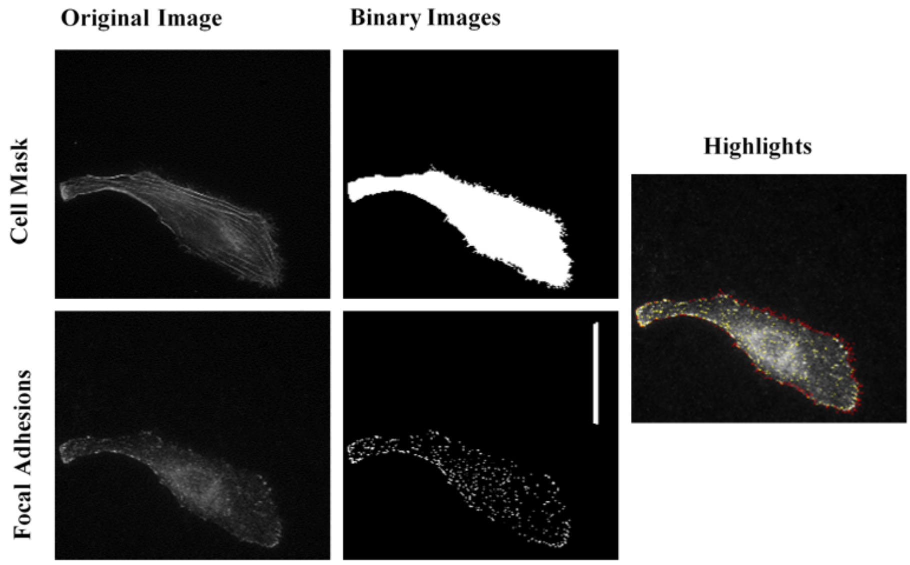


**Figure S10.** Steps of image processing for focal adhesion analysis. Scale bar: 50 µm.

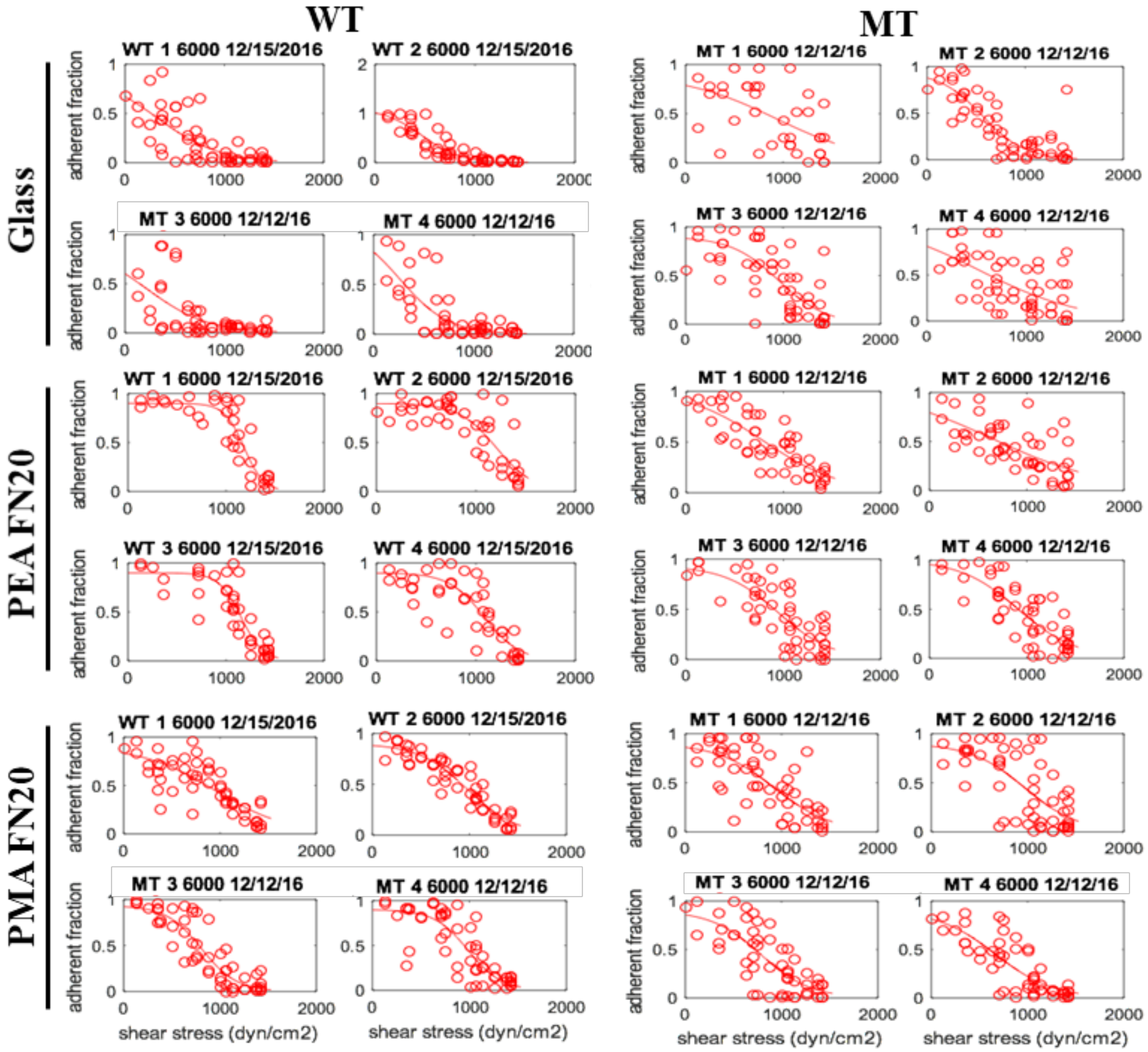

**Figure S11.** Cell adhesion strength measurements: Role of contractility. Detachment profile showing fraction of adherent cells versus applied shear stress for cells treated with BB adhering to substrates coated with FN solutions of 20 µg/mL for 2 h - FBS.

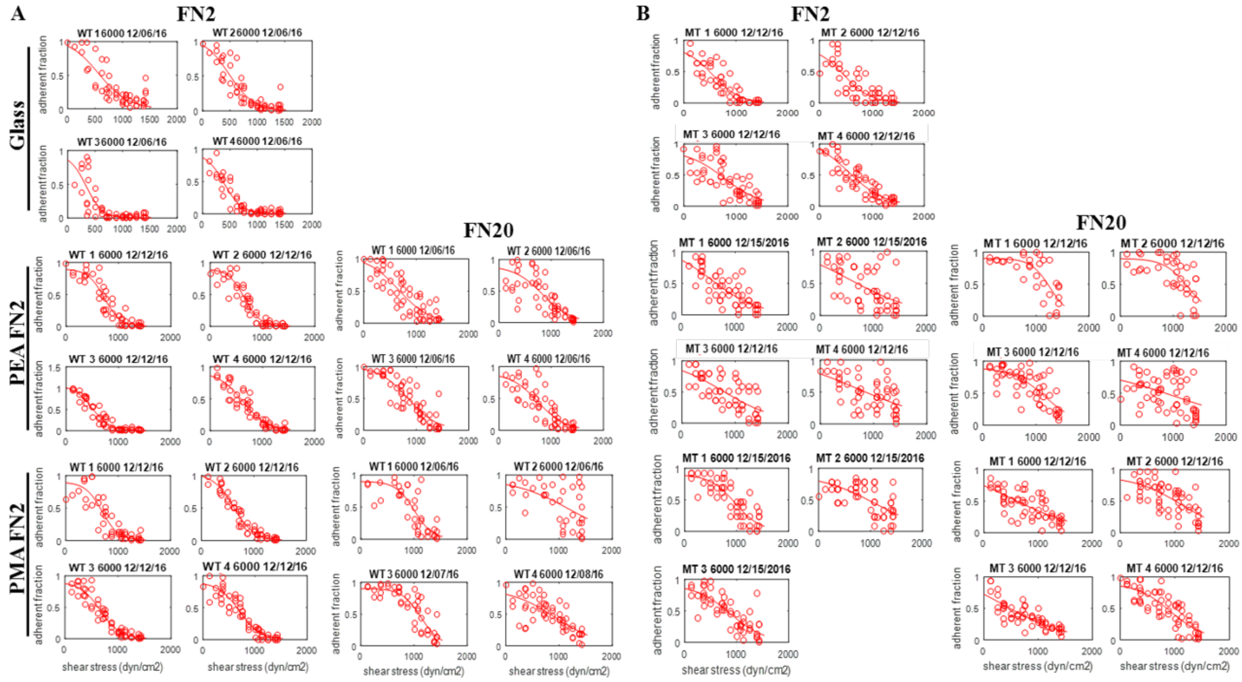


**Figure S12.** Cell adhesion strength measurements: Influence of fibronectin concentration. Detachment profile showing fraction of adherent cells versus applied shear stress for WT (A) and MT cells (B) adhering to substrates coated with FN solutions of 2 and 20 µg/mL for 2 h - FBS.

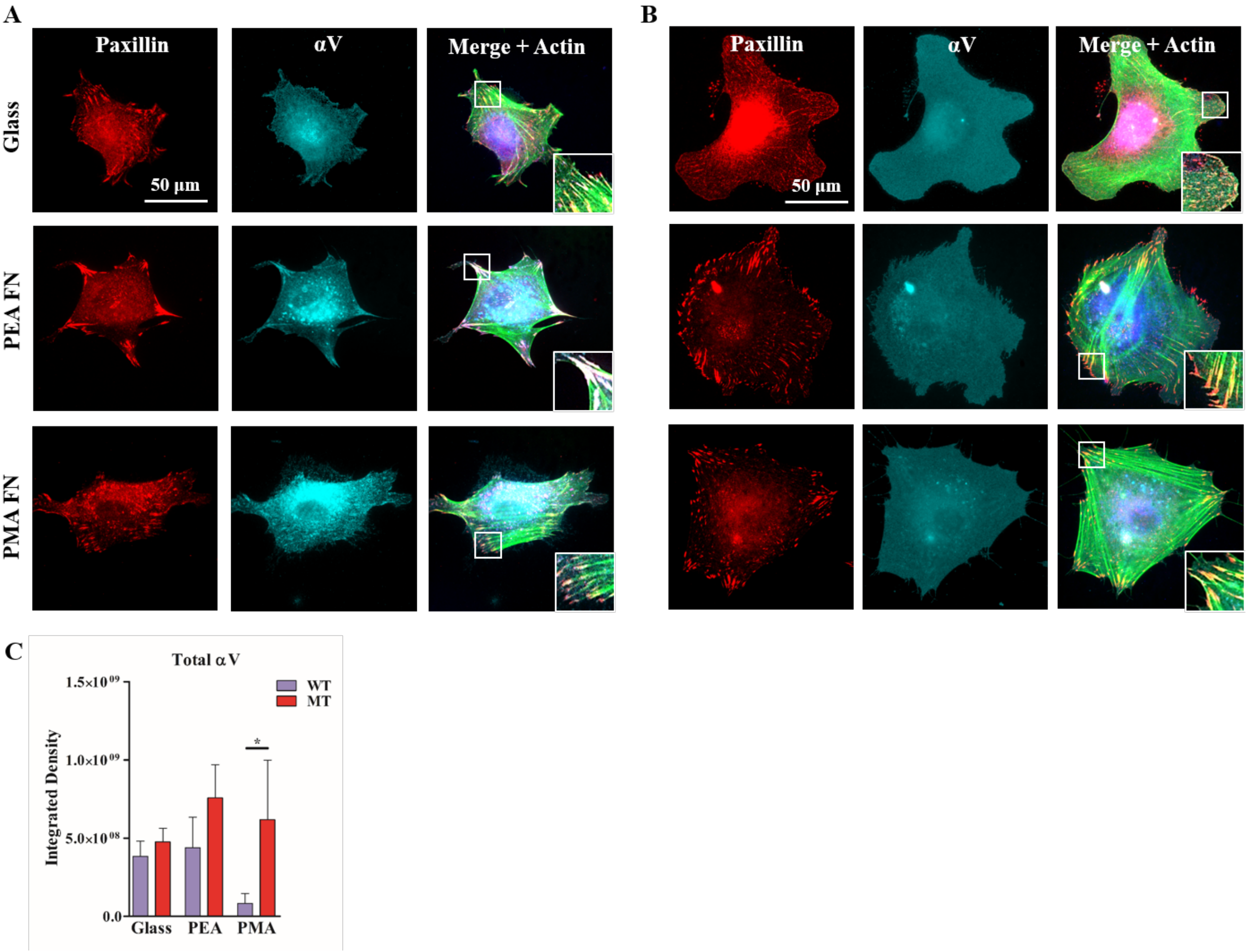


**Figure S13.** Integrin expression and focal adhesion by the MT and WT cells. Expression by WT (A) and MT (B) fibroblasts adhering to FN coated substrates for 2h under serum free conditions. Quantification of integrated density measurements of integrin αV (C) performed with ImageJ. Data are shown as the mean ± SD; n= 3. *$p < 0.05$. WT, wild type; MT, *COL4A2*$^{+/G702D}$ fibroblasts.

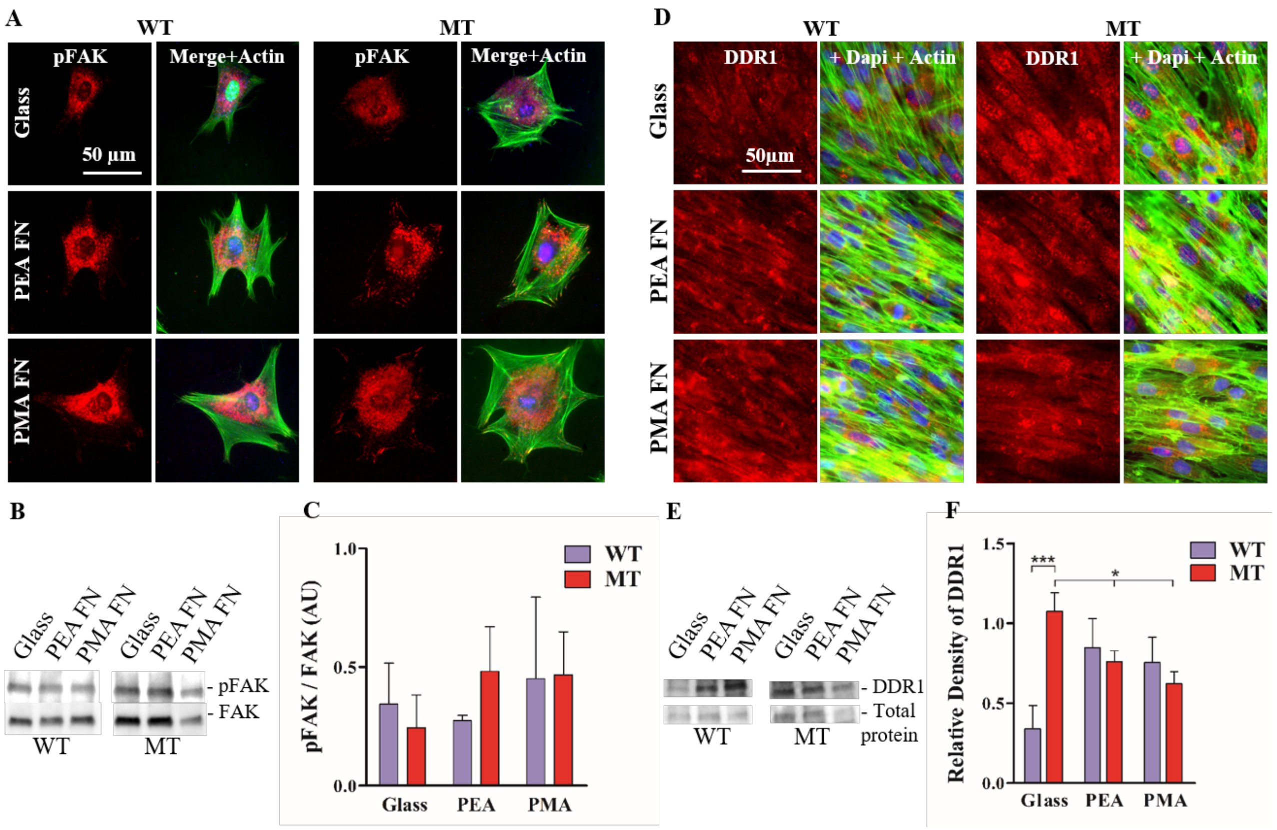


**Figure S14.** FAK phosphorylation and DDR1 analysis. Immunofluorescence images staining of phospho-FAK in WT and MT cells (A). Representative images of WB membranes with detected FAK and pFAK proteins in cells lysates after 2 h incubation (B); densitometry ratio of WB bands was calculated (C) n= 3. No statistically significant differences were found for the ratio. DDR1 (125 kDa) protein expression levels in the WT and MT cells assessed by IF (D) and western blot (E), the WB bands show traces of DDR1 in cells grown on FN coated substrates for 5 day. Total protein was used as the internal standard. Bands density was analyzed (F). Data are shown as the mean ± SD; n= 3. *p < 0.05; ***p<0.001. Immunofluorescence, IF. WT, wild type; MT, *COL4A2*$^{+/G702D}$ fibroblasts.

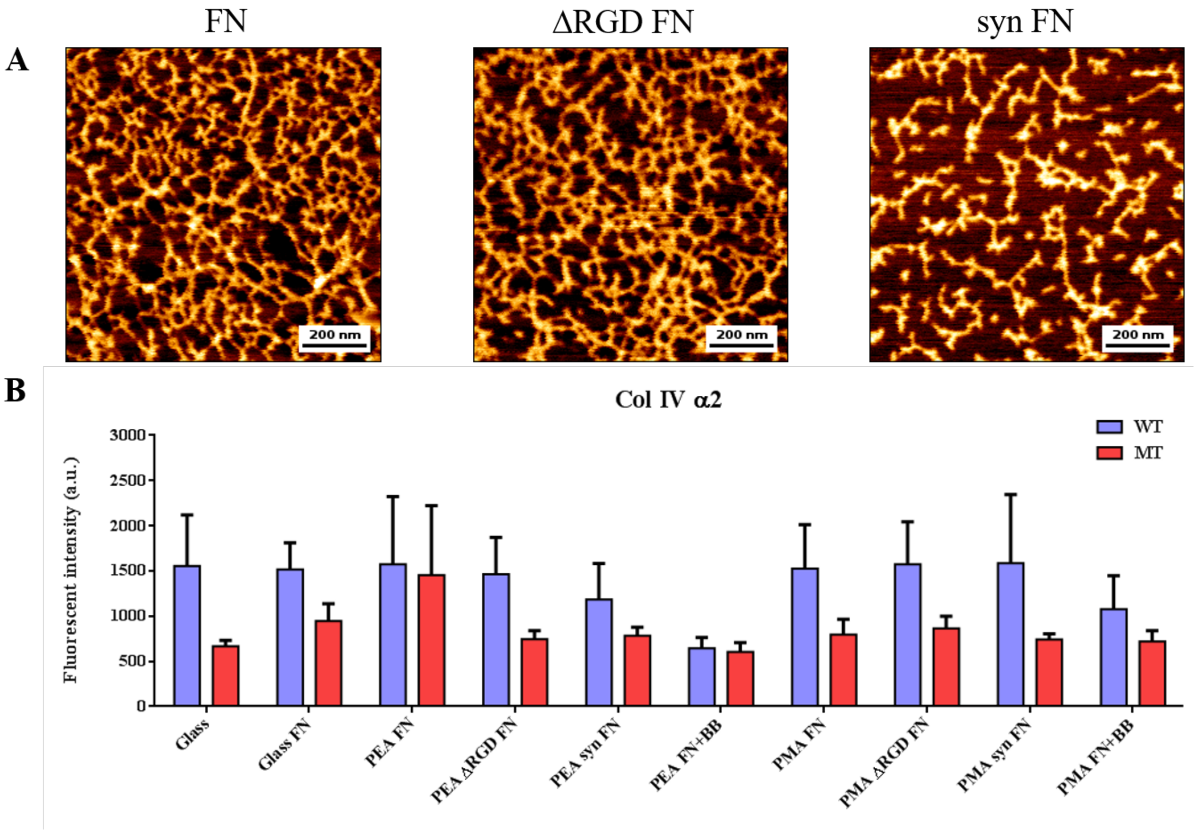


**Figure S15.** Secretion of Col4α2 by WT and MT fibroblasts on glass, PEA and PMA coated with mutant FN after 7 days of culture. (A) Mutant fibronectins formed interconnected nanonetworks upon adsorption on PEA, as seen via AFM phase imaging after coating of FN on PEA for 1 hour. FN is mouse wild type FN, ΔRGD FN is mouse FN lacking the RGD domain, syn FN is mouse FN with a mutation in the synergy binding site. (B) Quantification of integrated density of Col4α2 staining after variance filtering for cells grown on substrates (glass, PEA, PMA) either uncoated, coated with FN, ΔRGD FN, syn FN, or in the presence of blebbistatin (BB) in the culture medium.

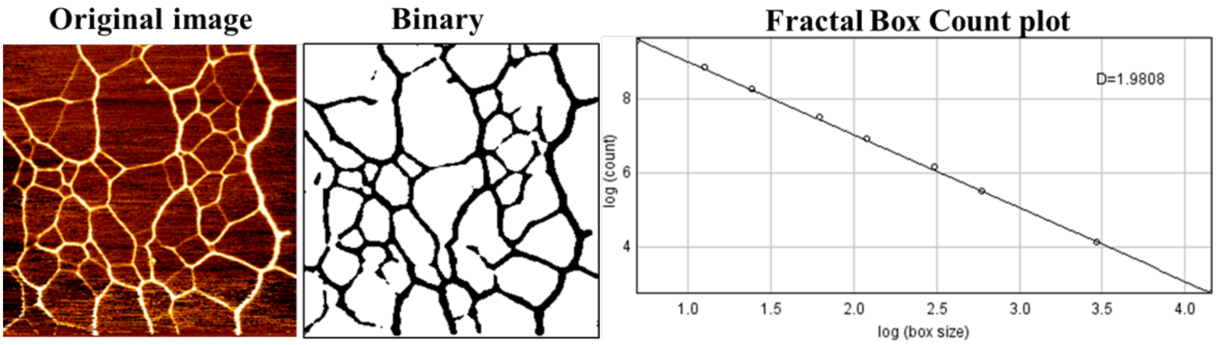


**Figure S16.** Image processing in order to prepare the image for the fractal box counting.
